# Gene Trajectory Inference for Single-cell Data by Optimal Transport Metrics

**DOI:** 10.1101/2022.07.08.499404

**Authors:** Rihao Qu, Xiuyuan Cheng, Esen Sefik, Jay S. Stanley, Boris Landa, Francesco Strino, Sarah Platt, James Garritano, Ian D. Odell, Ronald Coifman, Richard A. Flavell, Peggy Myung, Yuval Kluger

## Abstract

Single-cell RNA-sequencing has been widely used to investigate cell state transitions and gene dynamics of biological processes. Current strategies to infer the sequential dynamics of genes in a process typically rely on constructing cell pseudotime through cell trajectory inference. However, the presence of concurrent gene processes in the same group of cells and technical noise can obscure the true progression of the processes studied. To address this challenge, we present GeneTrajectory, an approach that identifies trajectories of genes rather than trajectories of cells. Specifically, optimal-transport distances are calculated between gene distributions across the cell-cell graph to extract gene programs and define their gene pseudotemporal order. Here, we demonstrate that GeneTrajectory accurately extracts progressive gene dynamics in myeloid lineage maturation. Moreover, we show that GeneTrajectory deconvolves key gene programs underlying mouse skin hair follicle dermal condensate differentiation that could not be resolved by cell trajectory approaches. GeneTrajectory facilitates discovery of gene programs that control the changes and activities of biological processes.

## 1 Introduction

Dynamic gene expression changes often specify mechanisms through which cells determine state and function. Indeed, tightly-regulated gene cascades underlie a myriad of fundamental processes, such as cell cycle/mitosis [1–4] and tissue/organ differentiation [5–8]. With the emergence of single-cell RNA-sequencing platforms, cell trajectory inference techniques [9–19] are widely applied to study the cellular dynamics of biological processes. These techniques use single-cell whole-transcriptome data to organize cells into lineages and infer a unidimensional latent variable (i.e., pseudotime [20]) that describes a cell’s position along a lineage process. After pseudotime construction, gene dynamics underlying a biological process can be inferred by tracking the changing patterns of their expression levels along the cell pseudotime [12, 15, 21].

However, when cells undergo multiple processes in parallel (e.g., cell cycle coupled with cell differentiation [22] or circadian clock [23]) and each process is governed by a different set of genes, cell pseudotime learned by organizing cells using the collective genes becomes less informative, as it mixes the effects of multiple processes. Mathematically, when multiple processes that are not strongly correlated with each other cooccur in the same group of cells, cell geometry (determined by these processes) cannot be effectively parametrized by a common single latent variable. Hence, organizing cells into unidimensional lineages is no longer appropriate.

To address this challenge, we propose GeneTrajectory, an approach to study dynamic processes that does not rely on unidimensional parameterization of the cell manifold. GeneTrajectory allows us to deconvolve multiple, independent processes with sequential gene dynamics. In contrast to cell trajectory approaches, GeneTrajectory constructs trajectories of genes rather than trajectories of cells. Our algorithm automatically dissects out gene programs from the whole transcriptome, eliminating the need for initial cell trajectory construction or the specification of the initial and terminal cell states for each process. Using this method, genes that sequentially contribute to a given biological process can be extracted and automatically organized into a respective gene trajectory that reveals the successive order of gene activity.

In this work, we begin by showing GeneTrajectory’s efficacy for unraveling gene dynamics through simulation experiments and application to a human myeloid lineage dataset. Subsequently, we employ our approach on a mouse embryonic skin dataset to demonstrate that GeneTrajectory can resolve critical cell state transitions during the early-stage development of hair follicles [5, 24]. Our results indicate that GeneTrajectory extracts gene geometry without the need for constructing cell pseudotime, revealing independent trajectories of concurrent processes that are otherwise obscured by cell pseudotime approaches.

## 2 Results

### 2.1 Computing optimal transport between genes over the cell graph

A progressive dynamic biological process is usually governed by a finely-regulated gene cascade [25–27], in which genes are activated and deactivated in a temporal order along the process, dictating the transcriptomic changes of underlying cell states. Moreover, cells can participate in multiple processes simultaneously, either in a dependent or independent manner. For instance, we illustrate two contrasting scenarios by considering the concurrence of a linear process (e.g. differentiation) and a cyclic process (e.g. cell cycle) (Fig. 1a). When these two process are strictly dependent on each other, they can be parameterized by a common latent variable and result in a 1-dimensional cell curve. In this scenario, it is straightforward to assign a meaningful pseudotime for the cells by ordering them along the curve. However, deconvolving genes into two processes and retrieving their pseudotemporal order in each process is not immediately apparent, which requires additional postprocessing (e.g., clustering gene dynamics along the cell pseudotime [12]). In contrast, when these two processes are independent, cells fall into a manifold (as a Cartesian product of these two processes) with an intrinsic dimension *>* 1. These processes do not share a common latent variable, thus gene dynamics inference based on unidimensional interpolation along the cell-cell manifold is no longer appropriate. In practice, the weak and stochastic nature of the dependency between concurrent biological processes can complicate the extraction of the cell path and the construction of cell pseudotime.

**Fig. 1.**
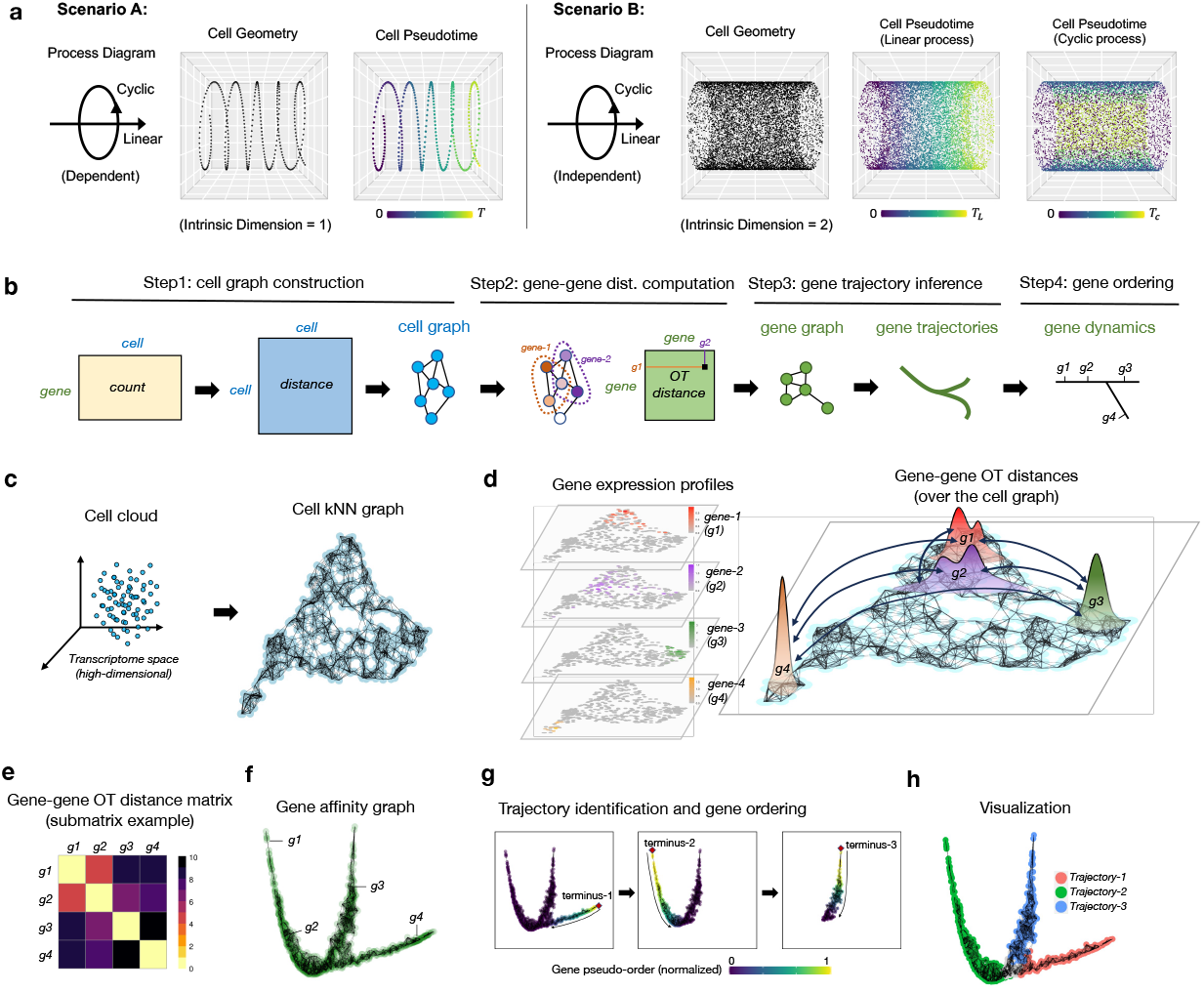
Overview of GeneTrajectory. a, Illustration of two scenarios when a linear process and a cyclic process are dependent or independent with each other, resulting in cell manifolds with different intrinsic dimensions and requiring distinct pseudotime parametrizations. b, Schematic of the major workflow of GeneTrajectory. c, Construction of cell kNN graph. d, Computation of graph-based OT (Wasserstein) distances between paired gene distributions (four representative genes are shown) over the cell graph. Gene distributions are defined by their normalized expression levels over cells. e, Heatmap of OT (Wasserstein) distances for genes g1-g4 in d. f, Construction of gene graph based on gene-gene affinities (transformed from gene-gene Wasserstein distances). g, Sequential identification of gene trajectories using a diffusion-based strategy. The initial node (terminus-1) is defined by the gene with the largest distance from the origin in the Diffusion Map embedding. A random-walk procedure is then employed on the gene graph to select the other genes that belong to the trajectory terminated at terminus-1. After retrieving genes for the first trajectory, we identify the terminus of the subsequent gene trajectory among the remaining genes and repeat the steps above. This is done iteratively until all detectable trajectories are extracted. h, Diffusion Map visualization of gene trajectories.

Here, we present GeneTrajectory, an approach to inferring gene processes through learning the gene-gene geometry without 1-dimensional parameterization of the cell manifold (Fig. 1b). Specifically, GeneTrajectory quantifies the distance of genes based on their expression distributions over a cell graph using optimal transport (OT) metrics (Fig. 1d). Previously, OT metrics (e.g., Wasserstein distance) have been applied in a wide range of scenarios in single-cell analysis, including 1) defining a distance measure between cells [28, 29] or cell populations [30], 2) constructing cell trajectories [31, 32], 3) spatial reconstruction of single-cell transcriptome profiles [33, 34], and multi-omics data integration [35]. In these works, the dissimilarity was quantified either between a pair of cells, or between a pair of cell populations. In our work, we distinctively define the graph-based Wasserstein distance between pairs of genes to study their underlying pseudotemporal dynamics. Specifically, we normalize the expression of a gene into a probabilistic distribution over cells and then compute the Wasserstein distances between gene distributions in the cell graph (Fig. 1d). Here, the cell graph is constructed in a way that provides a representation of cells which preserves the cell manifold structure in the high-dimensional space (Fig. 1c). In this construction, the graph-based Wasserstein distance between pairwise gene distributions has the following characteristics: 1) It takes into account the geometry of cells, i.e., it assigns a higher cost to transport a point mass from one cell to a distant cell as compared to its adjacent neighbors. 2) It prevents the transport across the ambient cell space which is often a problematic issue when using spatial distance measures (e.g., the Euclidean distance in the cell space).

In our approach, the computation of gene-gene Wasserstein distances is based on the following two steps:

- Construct a cell graph. As an initial step, we learn a reduced-dimensional cell embedding that can capture and represent the cell manifold structure in the original high-dimensional space. Next, we construct a *k*-nearest-neighbor (*k*NN) graph of cells based on their relative distances in the cell embedding (Fig. 1c). This establishes a cell-cell connectivity map that serves as the “roadmap” for transporting gene distributions in the next step. Here, for a given pair of cells *u* and *v*, we search for the shortest path connecting them in the *k*NN cell graph and denote its length as the graph distance (*d*_*G*_)_*uv*_ between cells *u* and *v*. This graph distance will be used to define the cost of transporting a point mass between cells *u* and *v* in the next step.
- Compute gene-gene Wasserstein distances over the cell graph. We model the expression level of genes as discrete distributions on the cell graph. Specifically, we divide the original expression level of a given gene in each cell by the sum of its expression level in all cells. We then define the distance between two gene distributions by the graph-based Wasserstein-p distance (*W*_*p*_-distance, 1 *≤ p < ∞*) (Fig. 1c,d). Accordingly, the transport cost between cell *u* and *v* is defined as 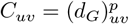. Here, *p* is a user-defined parameter, and *p* = 1 refers to the well-known Earth-Mover distance. Algorithmic details are described in Methods 8.1.2.

In practice, computing the Wasserstein distance between all pairwise gene distributions can be computationally expensive. When the cell graph is large, the time cost for finding the optimal transport solution increases exponentially. In our framework, we have designed two strategies to accelerate the computation based on 1) cell graph coarse-graining, and 2) gene graph sparsification (See details in Methods 8.1.2).

### 2.2 Gene trajectory construction

The gene-gene Wasserstein distance captures the pseudotemporal relations of genes in the sense that if two genes are activated consecutively along a biological process, their distributions are expected to have a substantial overlap in the cell graph and thus have a small Wasserstein distance between each other (Fig. 1e). To visualize the geometry of all genes, we convert pairwise gene-gene Wasserstein distances into genegene affinities and employ Diffusion Map to get a low-dimensional representation of genes. If dynamical cascades of gene activation and deactivation exist in the data, viewing the gene embedding by a combination of leading Diffusion Map eigenvectors delineates trajectories of genes (Fig. 1f). Each trajectory is linked with a specific gene program that dictates the underlying biological process.

In our approach, the extraction of gene trajectories is performed in a sequential manner (Fig. 1g). To identify the first trajectory, we search for the gene that has the largest distance from the origin of Diffusion Map embedding, which serves as the terminus of the first gene trajectory. To retrieve the other genes along the first trajectory, we take that terminus gene as the starting point of a diffusion process. Specifically, we assign a unit point mass to that gene and then diffuse the mass to the other genes. As the probability mass propagates along the gene trajectory from its terminus, the trajectory can be retrieved by a heuristic thresholding procedure (Methods 8.1.3). After retrieving genes for the first trajectory, we identify the terminus of the subsequent gene trajectory among the remaining genes and iterate the same procedure, until all detectable gene trajectories are extracted (Fig. 1g,h).

To order the genes along a given trajectory, we retain only these genes to recompute a Diffusion Map embedding based on their pairwise gene-gene Wasserstein distances. The obtained first non-trivial eigenvector of the Diffusion Map embedding provides an intrinsic ordering of the genes along that trajectory [36, 37].

To examine how the gene order along a given gene trajectory is reflected over the cell graph, we can track how these genes are expressed across different regions in the cell embedding. Specifically, we first group genes along each gene trajectory into successive bins and generate a cell embedding “snapshot” for each bin. In each snapshot, we color the cells according to the fraction of genes (from that bin) that they express. By plotting the expression level of each gene bin on the cell embedding, we can visualize how the underlying biological process progresses across cell populations.

### 2.3 Assessing GeneTrajectory’s performance using simulation

Assuming that a progressive biological process is temporally dictated by a sequence of genes, we simulated several artificial scRNA-seq datasets with a variety of gene dynamics by modeling the change of gene expression over time (Extended Data Fig. 1a,b, Methods 8.2.1). Specifically, for a gene involved in a given biological process, we simulate its expected expression level *λ*(*t*) as a function of time *t*. For clarity, we note that *t* represents the pseudotime of a biological process, linked with the cell state (e.g., differentiation status) rather than the actual time (e.g., specific day of a developmental process). Here, we use multiple parameters to account for the heterogeneity of gene expression profiles in single-cell data, including the variation of duration time and expression intensities (See details in Methods 8.2.1). For each cell state at *t* along a biological process, we apply a Poisson sampling to generate the observed expression level of each gene by taking *λ*(*t*) as the mean of Poisson distribution. In these simulation experiments, we know the ground truth of both the pseudotime of each cell in the corresponding biological process as well as the temporal order of genes that dictate each process. Lastly, we incorporate an optional step to account for sequencing depth. This is achieved by sampling a specified number of non-zero entries from the original count matrix. This procedure enables us to generate an artificial dataset with varying levels of missing data.

We first simulated 1) a cycling process in which the change of gene expression shows a periodical pattern over time (Fig. 2a, Extended Data Fig. 1c); and 2) a process with a branching point where it diverges into two different lineages (Fig. 2b, Extended Data Fig. 1d). Inspection of the gene trajectories in these two simulation examples reveals similar layouts with their cell embeddings (Fig. 2a,b). The ordering of genes along each gene trajectory shows a high concordance with the ground truth (Supplementary Table 1).

**Table 1.**
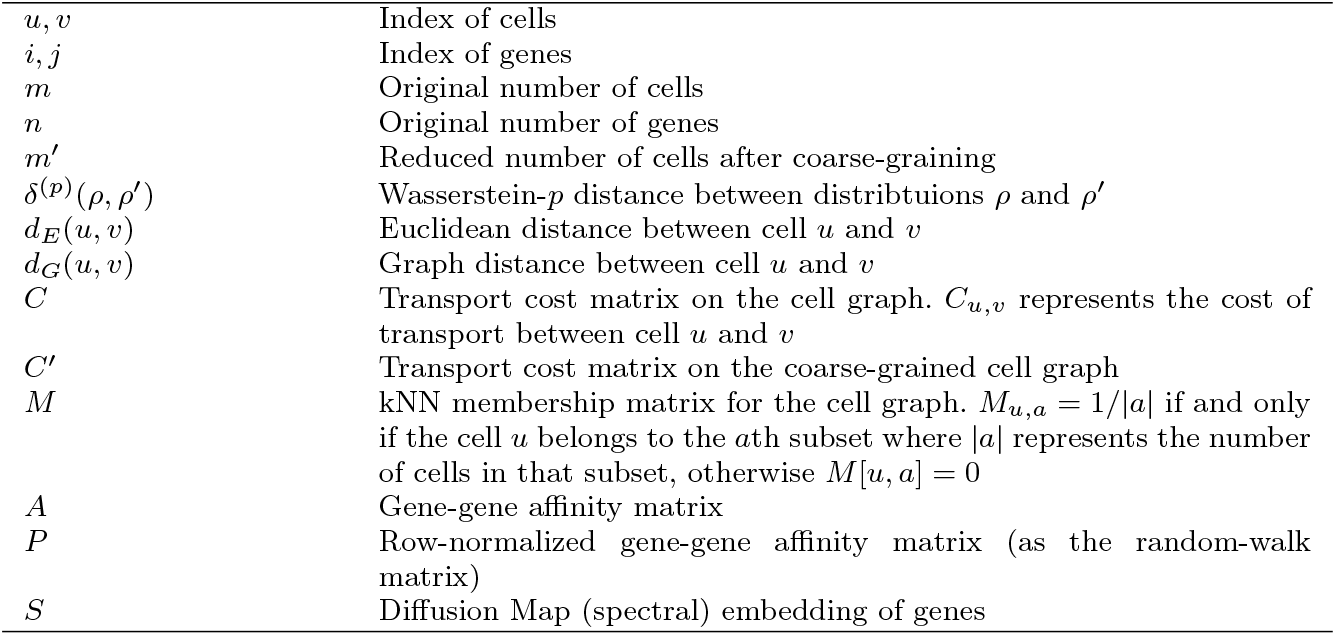
List of core notations in Methods.

**Fig. 2.**
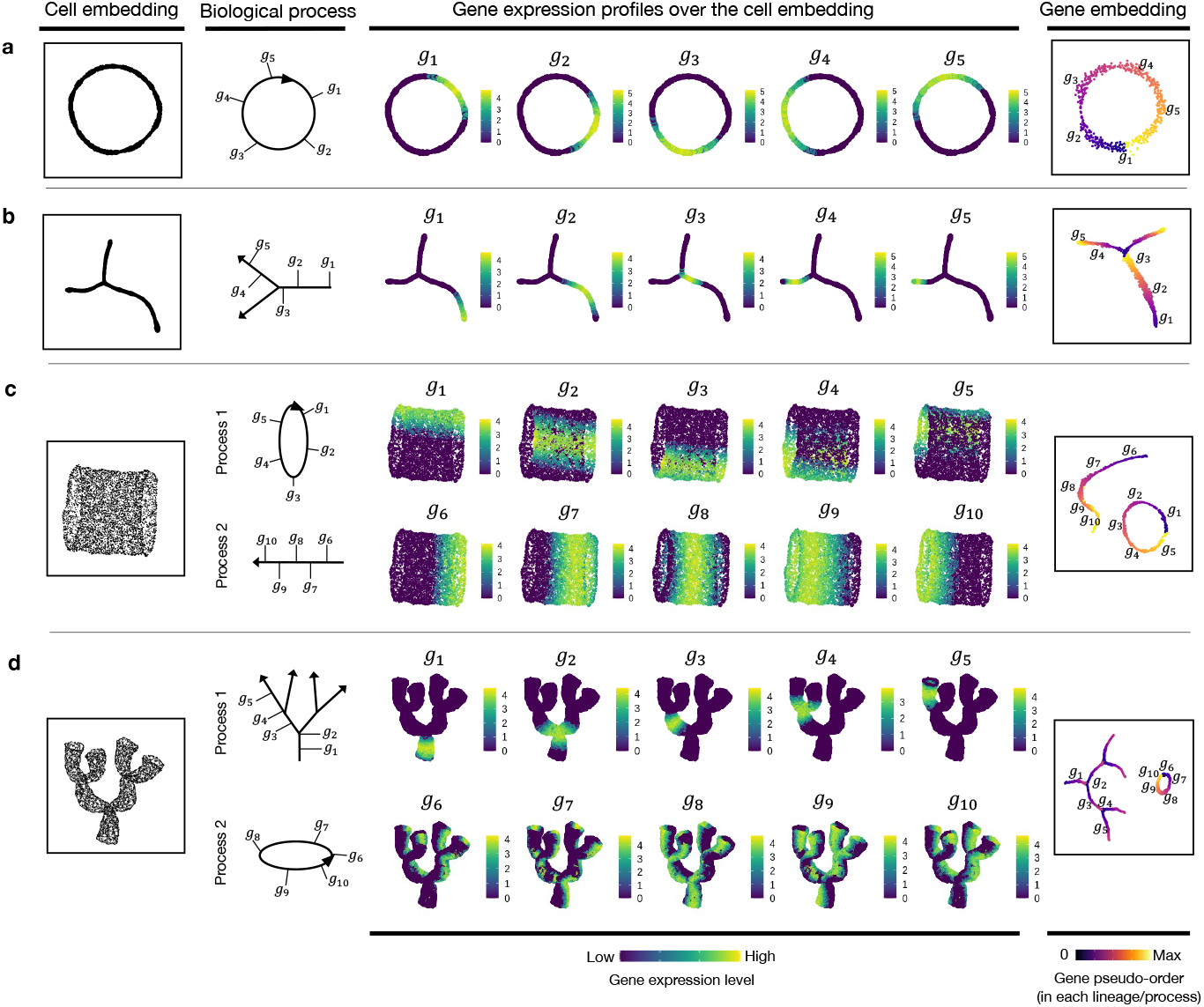
GeneTrajectory performance assessment based on simulation experiments. a, Simulation of a cycling process (cell cycle). The cell embedding and gene embedding showcase the same topology which has a ring-shaped structure. b, Simulation of a differentiation process with two lineages. The cell embedding and gene embedding showcase the same topology which has a bifurcating-tree structure. c, Simulation of a linear differentiation process coupled with cell cycle. The cell embedding and gene embedding showcase distinct topologies. Cells are organized along a cylinder-shaped manifold which has an intrinsic dimension of two. Genes that contribute to the two processes are deconvolved and organized along a ring-shaped trajectory and a linear trajectory. d, Simulation of a multi-level lineage differentiation process coupled with cell cycle. The cell embedding and gene embedding showcase distinct topologies. Cells are organized along a coral-shaped manifold which has an intrinsic dimension of two. Genes that contribute to the two processes are deconvolved and organized along a ring-shaped trajectory and a multilayered-tree-structured trajectory. [a and b are visualized by t-SNE; c and d are visualized by UMAP. The first column shows the cell embedding; The second column delineates the progressive dynamics of the simulated process with five genes selected along each process; The 3rd-7th columns show the expression of selected genes in the cell embedding following their pseudotemporal order; The 8th column displays the embedding of genes, colored by the ground truth of gene pseudo-order.]

We next, created two scenarios that simulate a mixture of two concurrent processes (Fig. 2c,d, Extended Data Fig. 1e,f). Specifically, one process mimics cell differentiation (linear or branched in a multilayered fashion), and the other mimics the cell cycle. In these two scenarios, each cell state is determined by two independent hidden variables – a pseudotime along the differentiation process and a pseudotime in the cell cycle. For each process, we simulated an exclusive set of genes with distinct dynamic characteristics (Extended Data Fig. 1e,f) (Methods 8.2.1), generating a cell manifold with a cylinder-shaped or a coral-shaped structure (Fig. 2c,d). In both scenarios, our approach deconvolves the original mixture of two processes into two gene trajectories representing a (linear or tree-like) differentiation process and a (circular) cell cycle process. Along each trajectory, genes are ordered in high concordance with the ground truth (Supplementary Table 1), indicating that GeneTrajectory allows deconvolving a mixture of biological processes that take place simultaneously in the same group of cells.

### 2.4 GeneTrajectory resolves myeloid gene dynamics

We demonstrate GeneTrajectory’s application using myeloid lineage differentiation, a classical biological system with a well-defined bifurcation of two major lineages [38, 39]. We extracted human myeloid cells from a public 10x PBMC dataset and identified four cell types based on canonical markers (Fig. 3a, Extended Data Fig. 2b-c). These included CD14+ monocytes, intermediate monocytes (HLA-DR high), CD16+ monocytes, and myeloid type-2 dendritic cells. The UMAP visualization of the cell embedding shows a continuum of cell states underlying myeloid lineage genesis, comprising monocyte maturation and dendritic cell differentiation. Human monocyte maturation involves upregulation of CD16 on a subset of CD14+ classical monocytes [40]. Specifically, CD14+ monocytes first transition into an intermediate subset of monocytes, and then differentiate into CD16+ non-conventional monocytes with distinct effector functions.

**Fig. 3.**
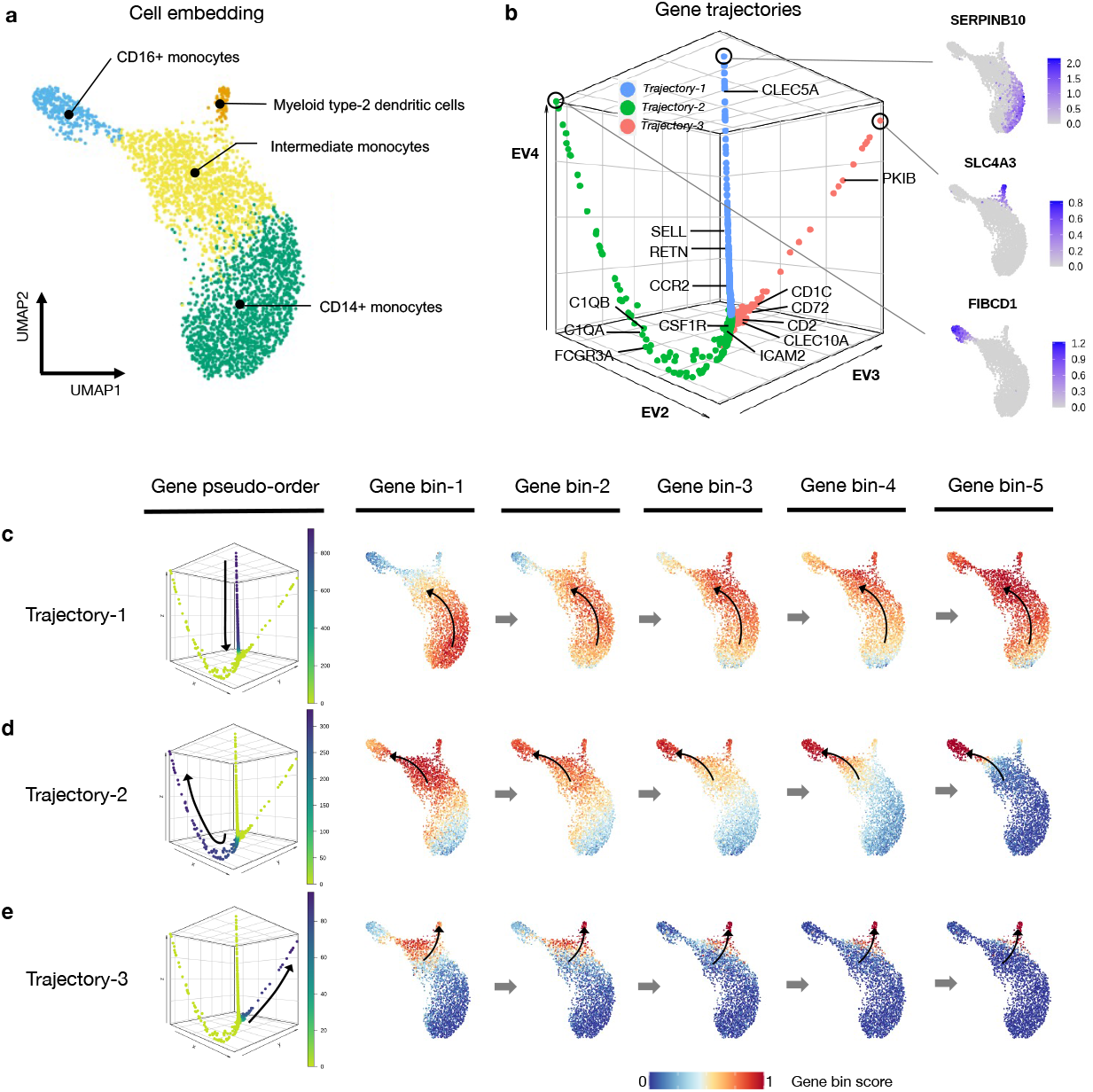
Gene trajectory inference on a myeloid scRNA-seq dataset. a, UMAP of myeloid cell population colored by cell types. b, DM embedding of the gene graph based on gene-gene Wasserstein distances, visualized using the three leading non-trivial eigenvectors. Three prominent gene trajectories are identified. Expression profiles of the genes at the terminus of these trajectories are shown, each indicating a distinct myeloid cell state. c-e, Gene bin plots showing the gene expression activities along each gene trajectory over the cell embedding. Genes along each trajectory are ordered and then split into five equal-sized bins. Gene bin score is defined by the proportion of genes (from each bin) expressed in each cell. Arrows indicate the path of gene distribution progression over the cells.

We used GeneTrajectory to identify three gene trajectories, each representing a specific aspect of myeloid lineage differentiation process (Fig. 3b). Viewing the gene bin plots of Trajectory-1 illustrates that a subset of CD14+ monocytes start a differentiation cascade and gradually shift towards CD16+ monocytes, which suggests Trajectory-1 captures the gene dynamics underlying the early stage of monocyte maturation (Fig. 3c). Notably, *CLEC5A, RETN, CCR2*, and *SELL* (CD62L) are known to be associated with the initial CD14+ monocyte cellular state [40] and are highlighted as part of Trajectory-1 (Fig. 3b). Subsequently, the ordering of genes that define Trajectory-2 provides a pseudotemporal view on the later stage of CD16+ monocyte differentiation (Fig. 3d). This process is primarily driven in response to cytokine CSF1 (colony-stimulating factor 1) and requires *CSF1R* [41]. While ordered after *CSF1R, ICAM2* is known to be constitutively expressed in CD16+ monocytes and is necessary for their patrolling ability across endothelium of blood vessels [41, 42]. Coming towards the end, *C1QA, C1QB* [43], *FCGR3A*, markers broadly expressed by fully differentiated CD16+ monocytes are identified. In addition, we retrieved a third gene trajectory (Trajectory-3) which marks the differentiation of type-2 dendritic cells as a distinct myeloid lineage (Fig. 3e). Myeloid type-2 dendritic cells have two subsets - CD14+ and CD14-. Specifically, the CD14+ subset shares overlapping features with CD14+ monocytes, whereas the CD14- subset is delineated here as corresponding with a separate gene trajectory [44]. In contrast to CD16+ monocytes, these CD14-dendritic cells differentiate in response to *GMCSF* and *IL4*, in line with expression of *CCR5, CD2, CLEC10A, CD72, CD1C, PKIB* [45] (Fig. 3b, Extended Data Fig. 2a). Notably, GeneTrajectory does not necessitate specification of the initial and terminal cell states for each process, while those states can be automatically revealed by inspecting the cell population that express the endpoint genes of each gene trajectory.

### 2.5 Deconvolving gene processes in dermal condensate genesis

Hair follicle dermal condensates (DCs) emerge in the skin dermis around embryonic day 14.5 (E14.5) and play an essential role in hair follicle formation. Morphogenetic signals, including Wnt/*β*-catenin signaling, are critical for the differentiation of DC cells [5, 24, 46]. We collected skin from E14.5 wildtype and paired K14Cre;Wntless^fl/fl^ (Wls) mutant embryos for single-cell RNA sequencing (Fig. 4a). The genetic defect in the mutant results in attenuated dermal Wnt signaling and a lack of DCs and hair follicles [47, 48] (Fig. 4b-c, Extended Data Fig. 3a).

**Fig. 4.**
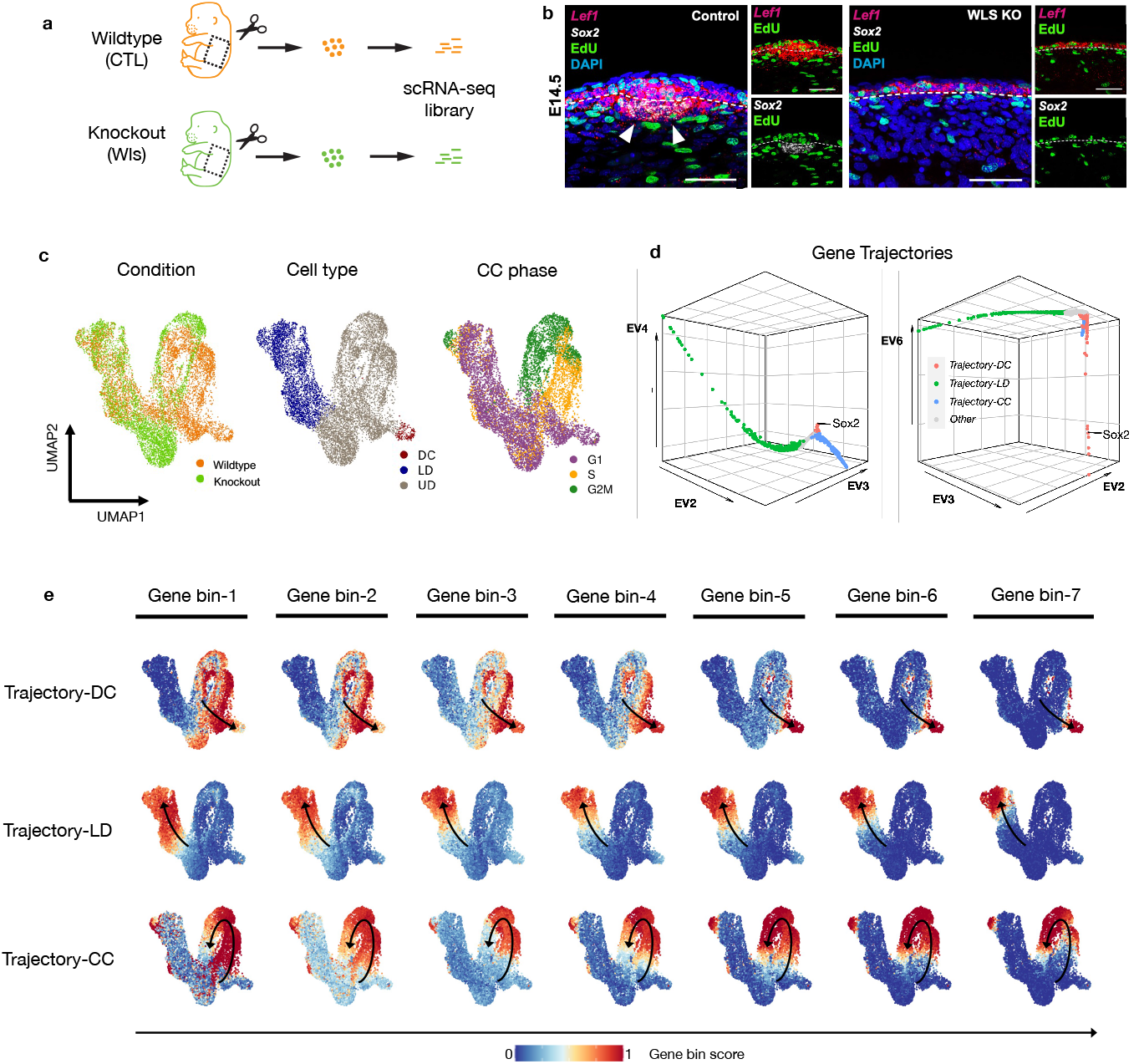
GeneTrajectory deconvolves two mixed processes during dermal condensate (DC) genesis. a, Experimental design of extracting skin tissue from a pair of wildtype and Wntless (Wls) knockout embryos at day E14.5 for single-cell RNA sequencing. b, Fluorescent in situ hybridization (FISH) images (scale bar = 50*µ*m) showing the spatial distribution of *Lef1, Sox2*, EdU nucleotide, and DAPI in the upper dermis of wildtype and Wls knockout. EdU is a nucleotide that is incorporated by cells in the S-phase of cell cycle. *n* = 8 (WT) and *n* = 9 (KO) embryos examined over 4 biologically independent experiments with similar results. c, UMAP of cells color-coded by cell types, conditions, and cell cycle phases. d, DM embedding of the gene graph to visualize three identified gene trajectories (two different combinations of leading non-trivial eigenvectors are displayed). e, Gene bin plots delineating the dynamics of each process (including DC differentiation, LD differentiation, cell cycle), in which genes along each trajectory are split into 7 equal-sized bins. Gene bin score is defined by the proportion of genes (from each bin) expressed in each cell. Arrows indicate the path of gene distribution progression over the cells.

Visualizing cells on UMAP reveals a continuum of cell states composed of lower dermal cells (*Dkk2* +) and Wnt-activated upper dermal cells (*Dkk1* + or *Lef1* +) which include DC cells (*Sox2* +) (Fig. 4c, Extended Data Fig. 3a). We applied GeneTrajectory to the combined dermal cell populations and extracted three prominent gene trajectories that correspond to lower dermis (LD) differentiation, dermal condensate (DC) differentiation, as well as cell cycle (CC) (Fig. 4d). Specifically, we examined the CC gene ordering by checking the distribution of genes associated with different CC phases along the gene trajectory (Extended Data Fig. 3b). Wnt signaling pathway genes (e.g., *Lef1, Dkk1*) and SHH-signaling pathway genes (e.g., *Ptch1, Gli1*), two morphogenetic signals shown to be necessary and sufficient for DC differentiation [5], are present in the DC gene trajectory. Notably, the upper dermal cell embedding integrates a mixture of biological processes (CC and DC differentiation) that co-occur within the same cell population. By employing GeneTrajectory, each biological process can be deconvolved from the other and independently examined. Viewing the gene bin plots for the CC and DC gene trajectories together reveals that DC progenitors actively proliferate throughout all stages and then exit the cell cycle at the terminus of DC differentiation (Fig. 4e). These data imply that DC cells are immediate progeny of proliferative progenitors in the upper dermis.

### 2.6 GeneTrajectory identifies biological defects in Wls mutant

We next use GeneTrajectory to examine how attenuated Wnt signaling affects the DC differentiation gene program. By tracking the expression status of genes along each gene trajectory in the wildtype and mutant cells (Fig. 5a), we did not detect a difference between the mutant and control with respect to the CC and LD gene trajectories (Extended Data Fig. 4a-b). However, along the DC gene trajectory, Wls mutant cells fail to express later-stage DC markers, indicating the defect is specific to DC differentiation. Visualizing gene bin plots for the DC gene trajectory show that mutant cells fail to progress in the DC differentiation process (Fig. 5e-f).

**Fig. 5.**
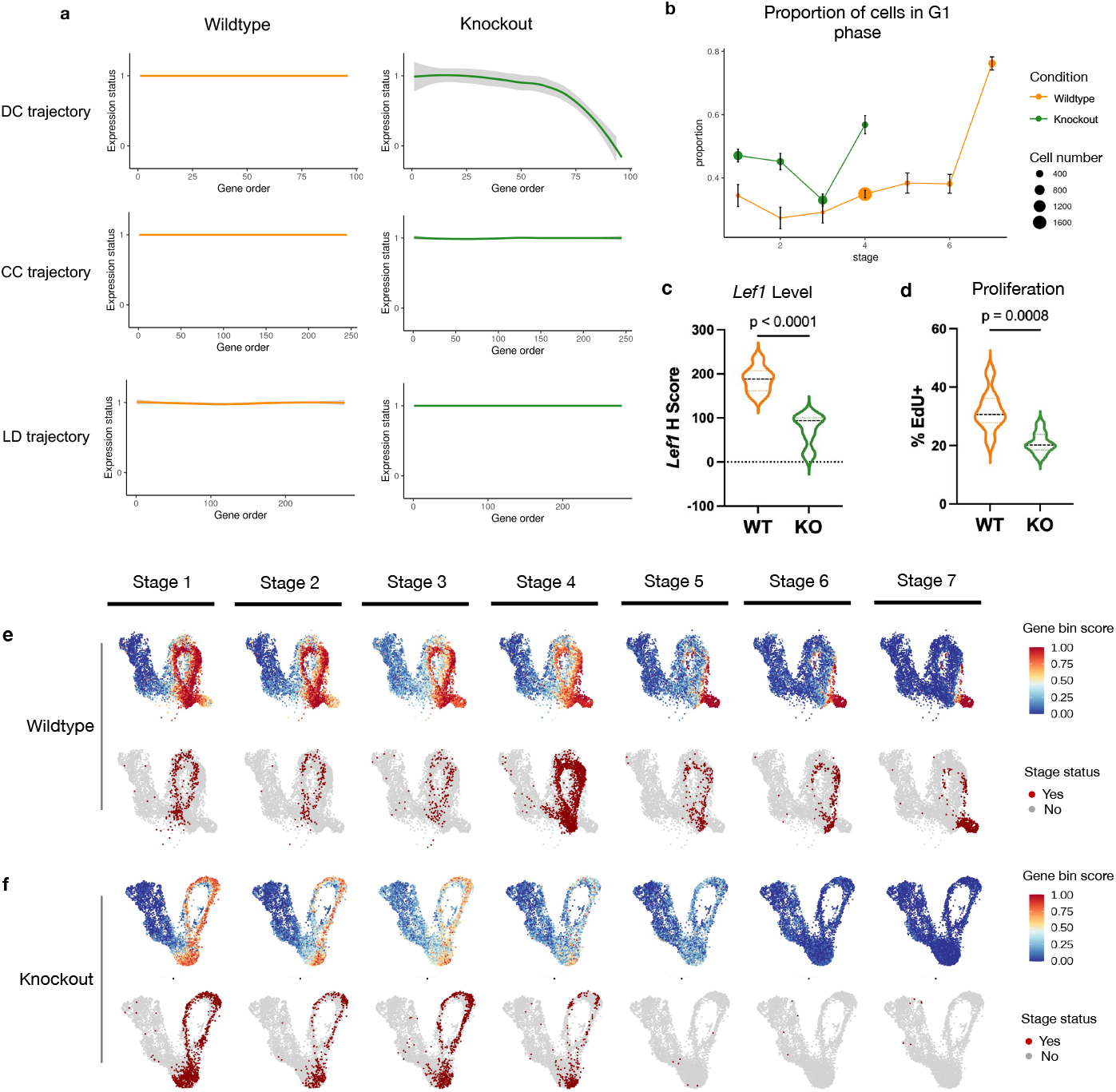
Gene dynamics comparative analysis. a, Gene expression status (smoothed) along each gene trajectory in two conditions (0: expressed in fewer than 1% of cells, 1: otherwise). b, Change in G1 proportion across seven stages of DC differentiation. Error bar: mean *±* SE, *n* = the number of cells in each stage of the corresponding condition. Dashed lines: the number of cells in the subsequent stage *≤* 10. c, *Lef1* transcript levels (H score) quantified by FISH in UD in two conditions. d, Percentage of Edu in UD in two conditions. EdU is a nucleotide incorporated by cells in the S-phase of CC. e, Gene bin plots of the DC gene trajectory over the wildtype cell embedding. Cells involved in the DC differentiation process are stratified into seven different stages. f, Gene bin plots of the DC gene trajectory over the Wls KO cell embedding. Cells involved in the DC differentiation process are stratified into seven different stages. Violin plots: *n* = 8 (WT) and *n* = 9 (KO) embryos examined over 4 biologically independent experiments. Statistical analysis was performed using two-sided student’s t-test. Lines indicate 75th, 50th and 25th percentiles.

Moreover, GeneTrajectory inference allows us to define a specific stage of cell state transition by specifying a gene window along the gene trajectory. To understand how the genetic mutation affects different stages of DC differentiation, we use GeneTrajectory to stratify the pool of progenitors by different stages of DC differentiation. Considering genes in each bin as markers indicative of a specific stage during DC differentiation, we first identified cells that express more than half of the genes in the last bin as cells in the final stage of differentiation (Stage 7). Among the remaining cells, we identified the cells that express more than half of the genes in the sixth bin as progenitors in Stage 6. We repeat this procedure iteratively until all seven gene bins were associated with their matched cell populations (Fig. 5e-f, Extended Data Fig. 4c).

By comparing the composition of progenitors in different stages between the wild-type and Wls mutant, we found that mutant cells fail to express most of the markers after Stage 4, when key markers in Wnt (e.g., *Lef1*) and SHH (e.g., *Gli1, Ptch1*) signaling pathways are upregulated in the wildtype condition (Fig. 5e-f, Supplementary Table 2). The average expression level of Wnt target genes is uniformly lower in the mutant than in the wildtype condition (Fig 5c, Extended Data Fig. 4d), while the proportion of cells in the G1 phase of the cell cycle is higher in the mutant across all stages (Fig. 5b). Consistent with this, the rate of EdU nucleotide incorporation (S phase) is lower in the Wls mutant (Fig. 4b, Fig. 5d). These data suggest that higher levels of Wnt signaling are necessary to maintain a normal rate of cell proliferation across the DC differentiation process until DC progenitors exit the cell cycle at the Stage 7. These results also raise the notion that dermal proliferation itself may directly regulate dermal cell state progression during the DC differentiation process.

### 2.7 Comparison of GeneTrajectory to cell trajectory methods

We compared GeneTrajectory with five cell trajectory methods, Monocle2 [16], Monocle3 [10], Slingshot [9], PAGA [11], and CellRank [15]. In the simulations of two co-occurring processes, we assessed performance by calculating the Spearman correlation between the gene ordering inferred from each approach and the ground truth. To order genes based on these cell trajectory inference methods, we first constructed the cell pseudotime using their default pipelines (Methods 8.2.6). Subsequently, we fitted Generalized Additive Models (GAM) models [49, 50] to find the peak location of each gene expression along the cell pseudotime. The genes were then ordered based on these peak locations. GeneTrajectory achieved the best performance in recovering gene order for both cyclic and linear processes (Fig. 6a,b) in simulation experiments, showing remarkable robustness to variations in cell numbers and sparsity levels of the count matrix.

**Fig. 6.**
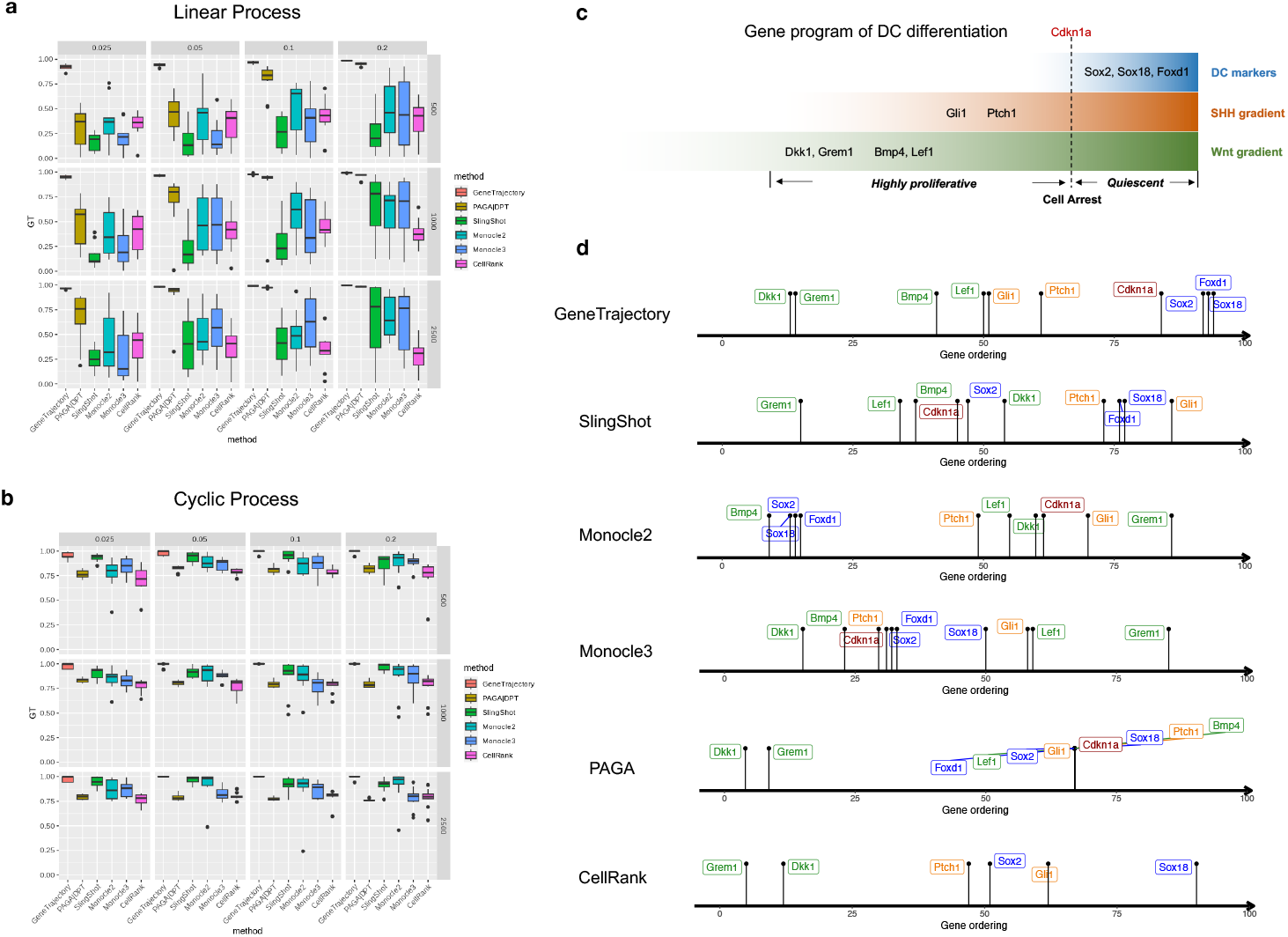
GeneTrajectory outperforms other methods in inferring gene ordering along concurrent processes. a-b, Comparison of GeneTrajectory with other approaches on simulated data of simultaneous linear and cyclic processes (corresponding to the third simulation example in Figure 2) with varying sample size and sparsity level of the count matrix (The numbers in the vertical gray boxes correspond to sample size, and those in the horizontal gray boxes correspond to the percentage of non-zero entries in each count matrix). c, Schematic representation of the key genes activated during the dermal condensate differentiation process. d, Gene ordering results obtained by different methods on the dermal condensate genesis data. Box plots: The box represents the interquartile range (IQR), with the line inside the box indicating the median. Whiskers extend to a maximum of 1.5 × IQR beyond the box, with outliers represented as individual points.

In our real-world example of DC development, we examined the order of known markers during DC differentiation (Fig. 6c,d). GeneTrajectory recovered the correct ordering: Wnt target genes *Dkk1/Grem1/Lef1* and *Bmp4* emerge first along this process. Dermal Wnt signaling is known to be required for SHH activation [47, 48]. Accordingly, the emergence of Wnt target genes is succeeded by the expression of SHH target genes (*Gli1/Ptch1*), which precedes the upregulation of the cell cycle inhibitor, *Cdkn1a*, and terminates with the expression of mature DC markers (*Sox2/Sox18/Foxd1*). In contrast, SlingShot, Monocle2, and Monocle3 were unsuccessful in generating a reasonable sequence for these genes. PAGA failed to generate a distinguishable ordering of later-stage markers. CellRank incorrectly placed the DC marker (*Sox2*) before *Gli1* and failed to define the ordering for *Bmp4/Lef1* and *Cdkn1a*.

Moreover, manually regressing out known coexisting biological effects (e.g., cell cycle) does not guarantee an accurate recovery of gene dynamics when using cell trajectory inference methods. For instance, in our dermal example, regressing out cell cycle effects resulted in persistent incorrect gene orderings for SlingShot, Monocle2, Monocle3, PAGA, and CellRank (Extended Data Fig. 5), suggesting that cell cycle regression is not sufficient to deconvolve the intertwined gene dynamics. This underscores the advantage of GeneTrajectory that it is capable of detecting and disentangling multiple gene programs when they are present.

## 3 Discussion

We developed GeneTrajectory, an approach for constructing gene trajectories where each trajectory comprises genes organized in a pseudotemporal order that characterizes the transcriptional dynamics of a specific biological process. GeneTrajectory uses optimal-transport-based gene-gene dissimilarity metrics. These metrics naturally leverage the underlying geometry of the cell-cell graph to reveal a coherent relation among genes that are involved in progressive processes. Importantly, GeneTrajectory bypasses the need for constructing cell pseudotime, which is a common requirement in existing methods. This renders it broadly applicable in scenarios where cells do not form into clear lineages.

It is worthwhile to note that cell trajectory inference and gene trajectory inference can complement each other to address different types of questions. Cell trajectory inference aims to define biological processes by lineages of cells, while gene trajectory inference associates each process with a sequence of genes. As demonstrated above, when cells participate in concurrent processes, cell trajectory inference may fail to deconvolve them. Similarly, when one gene participates in multiple biological processes, theoretically, it should be placed at the joint of gene trajectories. However, if that gene is expressed across many cells, it may have a small Wasserstein distance to genes that are homogeneously expressed (“uninformative genes”). As a result, it will be colocalized with uninformative genes in the gene embedding, causing difficulty for GeneTrajectory to distinguish them. Moreover, there are multiple aspects of our proposed algorithm that could be further refined. For instance, the branch identification procedure requires interactive optimization and might exhibit instability if the branches differ substantially in lengths and sizes. In addition, GeneTrajectory cannot automatically infer the directionality of progression along each trajectory. The directionality can be determined by checking whether the endpoint genes in each trajectory are initial stage markers or terminal stage markers of the corresponding process. Another important aspect is that the idea of utilizing the optimal transport distance between genes over cell-cell graphs could have other potential applications beyond the inference of gene programs and their dynamics. Intuitively, after we compute the gene-gene affinity matrix, we can then iteratively improve the organization of cells by an optimal transport distance between the cells over the gene-gene graph. This approach warrants further investigation from theoretical and practical perspectives.

In this work, we demonstrated the utility of GeneTrajectory to unravel gene dynamics using scRNA-seq data. However, our method can be generalized to other single-cell modalities, including but not limited to scATAC-seq [51], spatial transcriptomics [52], etc. Specifically, we anticipate GeneTrajectory can be applied to resolve biological processes using dual modalities [53] at the same time. For instance, we can quantify the pairwise distances between the distributions of gene expression and chromatin accessibility, which facilitates understanding the interplay between epigenetic dynamics and transcriptomic dynamics that underlie biological processes.

## 4 Acknowledgements

The authors thank Junchen Yang and Manolis Roulis for fruitful discussions. Y.K. and X.C. acknowledge support by National Institute of Health (NIH) grant R01GM131642. Y.K. also acknowledges support by NIH grant UM1DA051410, U54AG076043, U54AG079759, P50CA121974, and U01DA053628. X.C. is also partially supported by NSF DMS-2237842. P.M. is supported by the NIH (NIAMS) grant R01AR076420.

## 5 Author Contributions

R.Q., X.C. and Y.K. conceived the project and designed the framework. R.Q. implemented the method, performed data analysis, and wrote the manuscript. X.C. developed the computational methodology based on mathematical theories and contributed to writing. P.M. performed the experiments and interpreted the findings. Y.K., P.M., J.S.S. and E.S. contributed to writing and offered vital insights in improving the work. E.S., P.M., R.A.F. and I.D.O. contributed to the overall biological interpretation. B.L. and R.C. offered conceptual insights related to the theoretical framework. F.S. assisted in software implementation. S.P. assisted in experimental data analysis. J.G. assisted in writing.

## 6 Competing Interests

R.A.F. is an advisor to GlaxoSmithKline, Zai Lab and Ventus Therapeutics. F.S. is employed as director by PCMGF Limited. I.D.O. is the founder and president of Plythera, Inc and receives research funding from Ventus Therapeutics and SenTry.

## 8 Methods

### 8.1 Workflow

The major workflow of GeneTrajectory comprises the following four main steps:

- Step 1. Build a cell-cell *k*NN graph in which each cell is connected to its *k*-nearest neighbors. Find the shortest path connecting each pair of cells in the graph and denote its length as the graph distance between cells.
- Step 2. Compute pairwise graph-based Wasserstein distance between gene distributions, which quantifies the minimum cost of transporting the distribution of a given gene into the distribution of another gene in the cell graph.
- Step 3. Generate a low-dimensional representation of genes (using Diffusion Map by default) based on the gene-gene Wasserstein distance matrix. Identify gene trajectories in a sequential manner.
- Step 4. Determine the order of genes along each gene trajectory.

#### 8.1.1 Step 1. Construct a cell-cell graph and define graph distances

##### Data preprocessing

The data preprocessing contains the following steps:

i. Standard preprocessing of the count matrix (*m* cells, *n* genes),
ii. Dimension reduction.

##### Standard preprocessing

The original count matrix (cell-by-gene) is first preprocessed by employing the standard pipeline in single-cell analysis, including library normalization, top variable gene selection, and scaling.

##### Dimension reduction

Due to the low-rank nature of single-cell data, we run dimensionality reduction on the original count matrix to generate a low-dimensional representation of the cell geometry (cell embedding). Commonly used methods include PCA, t-SNE, UMAP, Diffusion Maps, etc. By default, we apply PCA for the initial step of dimensionality reduction and retain the leading *N* (typically around 30 *−* 100) principal components (PCs). Then we use Diffusion Map to generate a manifold-preserving low-dimensional representation of cells. Specifically, for a given pair of cells *u* and *v*, we calculate the Euclidean distance *d*_*E*_(*u, v*) between their coordinates of the leading *N* PCs. We then convert it into an affinity measure *a*(*u, v*) using the following Gaussian kernel with a local-adaptive bandwidth:

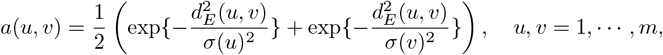

where *σ*(*u*) represents the Euclidean distance between cell *u* and its *k*-nearest neighbors in the PC space. Using a local-adaptive bandwidth allows to automatically adjust for the kernel size based on the local cell density in the original cell space. After we get the affinities between all pairs of cells, we apply the Diffusion Map algorithm and retain its leading *N* ′ eigenvectors as a low-dimensional representation of cells for the subsequent cell graph construction, which preserves the geometric information of the cell manifold.

##### Cell-cell graph distance computation

When cell geometry presents a low-dimensional manifold structure, the optimal transport should be always done across the cell manifold instead of taking a shortcut through empty regions in the ambient space where there are no cells. Here, we build a cell kNN graph in which we connect each cell to its *k*-nearest neighbors in the dimensionality-reduced cell space. For a given pair of cells *u* and *v*, we search for the shortest path connecting them in the kNN cell graph and denote its length as the graph distance *d*_*G*_(*u, v*) between cells *u* and *v*. Theoretically, in the limit of a large number of cells, the graph distances constructed in this way reveal manifold geodesic distances which are the intrinsic cell-cell distances [54, 55].

#### 8.1.2 Step 2. Compute graph-based Wasserstein distances between genes

We model the expression level of a gene as a discrete distribution on the cell graph. Specifically, let *g*_*i*_(*u*) represents the expression level of gene *i* in cell *u*, we then define the distribution of gene *i* by:

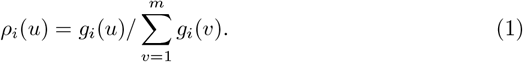

It has the following properties: 1)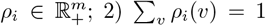. We then define the distance between two genes by the Wasserstein-*p* distance (*W*_*p*_-distance, 1 *≤ p < ∞*) between their distributions on the cell graph. Namely, the *W*_*p*_-distance *δ*^(*p*)^(*ρ*_*i*_, *ρ*_*j*_) between gene *i* and gene *j* quantifies their dissimilarity. Technically, the *W*_*p*_-distance can be computed by solving a discrete optimal transport mapping over the cell graph. Details are described below.

##### W_p_-distance formulation and computation

Here, we set up some mathematical notations: For a graph consisting of *m* nodes *V* = [*m*], a *graph distribution* is a non-negative vector 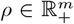 such that the sum of its elements is equal to one and the distribution assigns measure *ρ*(*u*) to node *u*. We assume the graph is equipped with a graph ground distance *d*_*G*_(*u, v*) for *u, v ∈ V*. Specifically, the graph distance *d*_*G*_ is used to specify the cost of the optimal transport, that is, the cost matrix *C* is defined as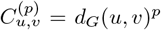. As mentioned in 8.1.1, we denote the shortest path distance on a kNN graph as *d*_*G*_, while the computational method also allows other options of *d*_*G*_ or even letting the cost matrix take a more general form. For two graph distributions *ρ* and *ρ*′, the *W*_*p*_-distance is defined as:

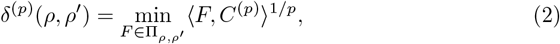

where Π_*ρ,ρ ′*_ = {*F, F*_*u,v*_ *≥* 0, Σ_*v*_*F*_*u,v*_ = *ρ*(*u*) for all *u*, Σ_*u*_ *F*_*u,v*_ = *ρ*′(*v*) for all *v*} denotes the set of transport plan *F* which pushes from the source distribution *ρ* to the target distribution *ρ*′.

##### Improve computational efficiency

In practice, the minimization in (2) can be solved by linear programming, which is computationally prohibitive on large cell graph and between all the pairs of genes. To reduce the cost of computing gene-gene *W*_*p*_-distances, we have designed two strategies to accelerate the computation based on 1) cell graph coarse-graining, and 2) gene graph sparsification. Briefly, cell graph coarse-graining aims to reduce the cell number by aggregating nearest cells into “meta-cells”. Gene graph sparsification aims to skip the computation for two gene distributions if they are very far away from each other at a coarse-grained level, as they are unlikely to participate in the same biological process. We note that while coarse-graining the cell graph to a crude scale can make it fast for computation, it may lose accuracy and compromise the resolution. Hence users should judiciously choose the level of coarse graining based on the capacity of their computing resources.

##### 1) Cell graph coarse-graining

We coarse-grain the cell graph by aggregating *m* cells into *m*′ “meta-cells” using K-means clustering algorithm. Specifically, let *M* be the *m*-by-*m*′ membership matrix where *M* [*u, a*] = 1*/* |*a*| if and only if the cell *u* belongs to the *a*th subset where |*a*| represents the number of cells in that subset, otherwise *M* [*u, a*] = 0, then we define an updated transport cost matrix *C*′ on the coarsegrained cell graph by *M*^*T*^ *CM*. Accordingly, the expression level of a given gene in each “meta-cell” is defined by the sum of its expression level in all the cells in that subset. Intuitively, this procedure can be viewed as providing an approximation of cell graph with fewer cell nodes.

##### 2) Gene affinity graph sparsification

We sparsify the gene affinity graph by zeroing out the entries where their pairwise Wasserstein distances are greater than a threshold.

The threshold is selected such that affinities associated with distances greater than it will be exponentially small and thus contribute negligibly to the gene affinity graph. The threshold is adaptively estimated for each cell using the approximate Wasserstein distance on a coarse-grained cell graph (Strategy 1) which allows fast computation.

Specifically, this is formulated in the following way: if we want to construct the gene-gene Wasserstein distance matrix on a cell graph of an original size *m*, we first coarse-grain *m* cells into *m*′ “meta-cells” using the procedure in Strategy 1 wherec *m*′ is a size that can be quickly handled. Based on the gene-by-gene Wasserstein distance matrix constructed on *m*′ “meta-cells”, we identify the *αk* nearest neighbors for each gene (where *α* is the predefined parameter and *k* is the neighborhood size to construct the local adaptive kernel for computing the Diffusion Map). Going back to the computation on the original cell graph, we then only compute the Wasserstein distance between a pair of genes if one of them is included in the other’s *αk* nearest neighbors. Practically, this can reduce the running time to 2*αk/m* of the original which computes Wasserstein distances for all pairs of genes.

#### 8.1.3 Step 3. Construct gene trajectories

After we obtain the gene-gene Wasserstein distance matrix, we convert it into an affinity matrix *A* using a local-adaptive Gaussian kernel. Specifically, the kernel bandwidth for each gene is defined by the distance to its *k*-nearest neighbor (as similar with 8.1.1). The affinity between gene *i* and gene *j* is defined by:

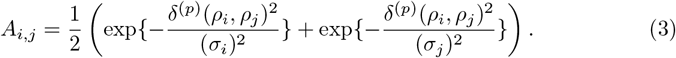

Here, *ρ*_*i*_ represents the distribution of gene *i*; *σ*_*i*_ represents the *k*-th smallest Wasserstein distance between gene *i* and other genes. *k* is an integer parameter to be specified by the user, which controls the size of the local neighborhood on the graph (in the sense that *A*_*ij*_ is only large on a subject of genes *j* which are sufficiently close to *i*). The affinity matrix *A* in (3) is used to construct a random walk on the gene-gene graph (see below in the bullet point “diffusion of probability mass on the gene graph”). The random walk constructed from affinity *A* allows us to apply Diffusion Map to obtain a low-dimensional embedding of the genes.

Next, extracting gene trajectories is processed in a sequential manner when the gene graph exhibits a tree structure. Briefly, we first identify an “extremum” gene as the terminus for the first gene trajectory, and then employ a diffusion strategy to retrieve genes belonging to that trajectory where the terminus gene servers as the initial node of the diffusion process.

The details are summarized below:

##### Selection of the initial node

We retain the top *d* nontrivial Diffusion Map eigenvectors as the low-dimensional spectral embedding of genes, denoted by *S*. Let *S*_*i*_ represents the spectral coordinates of gene *i*, we choose the gene with the largest *L*_2_ embedding norm 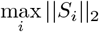 as the starting point of a diffusion on the gene graph. The assumption here is that the gene with the largest distance from the origin of spectral embedding corresponds to the terminus of a specific gene trajectory.

##### Diffusion of probability mass on the gene graph

The diffusion is performed by propagating a point mass from the initial node in the gene graph. Here, the initial probability mass *p*_0_ can be formulated as the following unit vector:

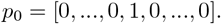

Suppose gene *j* is selected as the initial node, then only the *j*-th entry of *p*_0_ is equal to 1 while all other entries are zeros. We then construct a random-walk matrix *P* by row-wise normalizing the gene-gene affinity matrix *A*. Specifically, *P* is defined by:

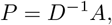

where *D* is the degree matrix of *A* (i.e., *D* is a diagonal matrix where *D*_*ii*_ = Σ_*j*_*A*_*ij*_). Calculating *p*_1_ = *Pp*_0_ gives the updated probability mass (over genes) after the first time of diffusion. We run the diffusion up to *t* times (the integer *t* is a tunable parameter) on the gene graph to get the *t*-step probability mass *p*_*t*_ = *P*^*t*^*p*_0_. We then select the genes {*j, s*.*t*., *p*_*t*_(*j*) *> τ*_0_ max_*j′*_*p*_*t*_(*j*′)} as members of the first gene trajectory. Here, *τ*_0_ is a thresholding parameter, which in practice can be set to be in the range of 0.02 *−* 0.05. Throughout the experiments in this paper, we choose *τ*_0_ = 0.02.

After the genes that belong to the first gene trajectory are extracted, we repeat the above procedure on the remaining genes to get the second gene trajectory, and then the third, etc. This algorithm allows retrieving a series of gene trajectories successively until all detectable trajectories are identified.

#### 8.1.4 Step 4. Order genes along each trajectory

To determine the gene ordering along a given gene trajectory, we first extract the corresponding sub-matrix of gene-by-gene Wasserstein distances as computed in 8.1.2. That is, we only focus on the genes which are the members of that trajectory. We then re-compute the Diffusion Map on the Wasserstein distance sub-matrix to obtain a new spectral embedding of genes in that trajectory. The first non-trivial eigenvector (EV2) of the new Diffusion Map embedding provides an ordering of the genes along that trajectory, according to the spectral convergence theory of Diffusion Map [36, 37]. Specifically, genes are ordered based on ranking their coordinates along EV2.

### 8.2 Experiments and analyses

Here, we present the details for the simulation experiments (8.2.1, 8.2.2), the biological experiments of mouse embryo skin sample preparation (8.2.3), the analyses on real-world biological datasets (8.2.4), comparing Wasserstein distance with other canonical metrics for learning gene geometry (8.2.5), comparing Gene Trajectory with cell trajectory methods in terms of gene ordering inference (8.2.6), and the robustness evaluation experiments and guidelines on parameter selection (8.2.7).

#### 8.2.1 Workflow of gene dynamics simulation

We present the details of our simulation framework for the four examples in Figure-2, including

A. a cyclic process,
B. a differential process with two lineages,
C. a linear differentiation process coupled with cell cycle,
D. a multi-level lineage differentiation process coupled with cell cycle.

For illustrative purposes, we first introduce the simulation procedure on a simple linear process. The corresponding plots are shown in Extended Data Figure 1a-b.

##### Illustrative example: a linear process (Extended Data Figure 1a-b)

As the simplest scenario, we simulate a linearly progressive biological process in [0, *T*] where *t* = 0 corresponds to the initial cell state and *t* = *T* corresponds to the terminal cell state. We simulate a set of genes {*g*_*i*_, *i* = 1, …, *n*}, where each *g*_*i*_ is a non-negative vector in ℝ^*m*^and *g*_*i*_(*u*) represents the gene expression at cell *u, u* = 1, …, *m*.

In this example, we let each cell *u* be uniquely associated with a pseudotime *t*_*u*_, which is i.i.d. uniformly distributed on [0, *T*]. Our procedure is to first construct for each gene *i* a continuous function *λ*_*i*_(*t*) on *t* ∈ [0, *T*], and then obtain the gene expression vectors *g*_*i*_ from *λ*_*i*_(*t*) based on Poisson sampling. Specifically, the simulation procedure consists of two steps:

##### Simulate expected gene expression levels along the process

For each *i*, we define a function *λ*_*i*_(*t*) where *λ*_*i*_(*t*_*u*_) represents the expected gene expression level of gene *i* at cell *u*. The function *λ*_*i*_(*t*) is associated with a “peak time” 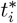, which represents the time point when gene *i* reaches the peak of its expected expression level. The time 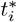 is uniformly sampled from [0, *T*]. The function *λ*_*i*_(*t*) then takes a parametric expression as

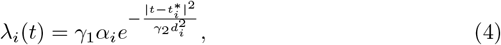

where parameters *γ*_1_, *γ*_2_ are predefined positive scalars, and *α*_*i*_, *d*_*i*_ are positive random variables to account for the variation in duration length and expression intensity of different genes. Specifically, we draw *d*_*i*_ and *α*_*i*_ from log-normal prior distributions as below:

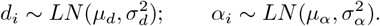

##### Sample gene reads from a Poisson distribution

In reality, the sequencing process is based on capturing molecules (e.g., DNA or RNA fragments) in a random manner. To mimic the randomness in the sequencing process, we simulate *g*_*i*_(*u*) as from a Poisson distribution with a rate *λ*_*i*_(*t*_*u*_), namely,

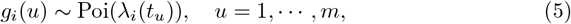

independently across all *u* and *i*. This gives 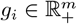 as desired.

##### (Optional) Sparsify the count matrix by sampling non-zero entries

Lastly, we incorporate an optional step to account for sequencing depth. This is achieved by randomly selecting a specified number of entries from the original count matrix without replacement, and subsequently zeroing out the remaining entries. The probability that an entry is selected is proportional to its original expression value. This procedure enables us to generate an artificial dataset with varying levels of missing data.

##### Example A: a cyclic process (Figure 2a)

To simulate a biological process with cyclic dynamics (e.g., cell cycle), based on the former setup of the linear process simulation, we only need to modify 4 as:

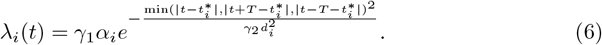

All other details are the same with 8.2.1.

##### Example B: a differentiation process with two lineages (Figure 2b)

To simulate a biological process with a bifurcating structure (e.g., myeloid lineage differentiation), we represent the underlying cell state by a generalized pseudotime vector *τ* comprising three pseudotime variables:

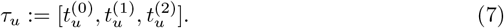

Here,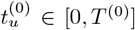 represents the pseudotime along the initial process prior to lineage differentiation, 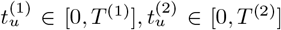 each represents the pseudotime along the process of lineage-1, lineage-2 differentiation.

Specifically, if a cell *u* is along the initial process, then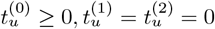. If a cell *u* is along lineage-1, then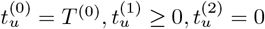. If a cell *u* is along lineage-2, then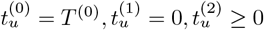.

Similarly, we generate 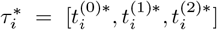 based on the same procedure as described above to represent the “time point” that gene *i* reaches the peak of its expected expression level. Here, the expectation of the expression level of gene *i* at the time point *τ* is given by:

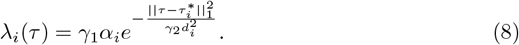

Parameters *γ*_1_, *γ*_2_, *α*_*i*_, *d*_*i*_ are defined based on the same procedure in 8.2.1. We then simulate

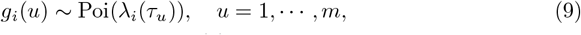

independently across *u* and *i*, similarly as in (5).

##### Example C: a linear differentiation process coupled with cell cycle (Figure 2c)

In this example, we simulate genes for a linear process and a cyclic process independently and then put them together. Specifically, we associate each cell *u* with a generalized pseudotime vector *τ* comprising two pseudotime variables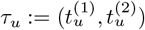.

Here, 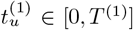represents the pseudotime along the linear process, while 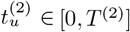 represents the pseudotime along the cyclic process. The sampling processes to generate 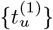 and 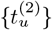 are independent.

Next, we simulate two sets of genes using the procedure same as in 8.2.1 but with a different definition of the Poisson rate function *λ*_*i*_(*𝒯*). Specifically, the first list of genes {*g*_*i*_}, 1 *≤ i ≤ n*_1_ are simulated with

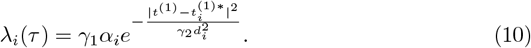

The second list of genes {*g*_*j*_}, *n*_1_ *< j ≤ n* are defined with

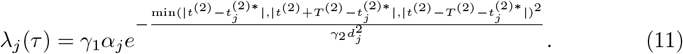

Notably, the first list of genes contribute to the linear process while the second list contribute to the cyclic process. This simulation results in a cylinder-like cell manifold in the high-dimensional space.

##### Example D: a multi-level lineage differentiation process coupled with cell cycle (Figure 2d)

In this example, we simulate genes for a 2-level tree-structured process and a cyclic process independently and then put them together. In the general case, let us consider simulating a *N* -level bifurcating process, in which the initial process *P*_0_ first splits into two lineages (*P*_1_ and *P*_2_), then each lineage proceeds independently and further splits into another two sub-lineages (*P*_1.1_ and *P*_1.2_, *P*_2.1_ and *P*_2.2_), and each sub-lineage divides again in an iterative manner until a *N* -level tree structure is generated. At the same time, all the cells are involved in a cyclic process. Here, each cell *u* can be associated with a generalized pseudotime vector *𝒯* comprising 2^*N*^ pseudotime variables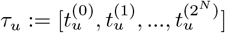, each of the first 2^*N*^ *−* 1 pseudotime variables corresponds to a pseudotime location along *P*_0_, *P*_1_, *P*_2_, *P*_1.1_, *P*_1.2_, *P*_2.1_, *P*_2.2_, …, 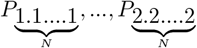 respectively. For generating the instances of these 2^*N*^ − 1 pseudotime variables, we adopt the similar framework as applied in 8.2.1 by requiring that when a cell *u* is along a daughter lineage, its pseudotime variables corresponding to all the parent processes are set to the largest possible values, and its pseudotime variables corresponding to other processes (exluding the daughter lineage itself) are all set to 0. Besides, 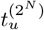 represents the pseudotime of cell *u* in the cyclic process, which is independent from the other 2^*N*^ − 1 pseudotime variables.

Next, we simulate two sets of genes using the procedure same as in 8.2.1 but with a different definition of the Poisson rate function *λ*_*i*_(*τ*). Specifically, the first list of genes {*g*_*i*_}, 1 *≤ i ≤ n*_1_ are simulated with

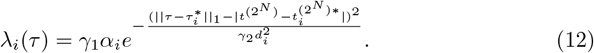

The second list of genes {*g*_*j*_}, *n*_1_ *< j ≤ n* are defined with

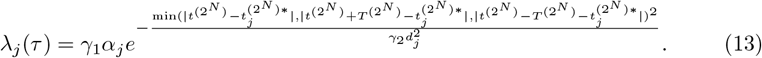

Notably, the first list of genes contribute to the tree-structured differentiation process while the second list contribute to the cyclic process. This simulation results in a coral-like cell manifold in the high-dimensional space.

##### Details of the simulation examples shown in Figure 2

For the examples shown in Figure 2, the evaluation outputs can be found in the Supplementary Table 1. Each example was tested through 10 replicates. Specifically, in the first example, we simulated 1, 000 cells, 500 genes underlying the cyclic process. In the second example, we simulated 1, 000 cells, 500 genes for the initial process and 250 genes for each daughter lineage process. In the third example, we simulated 5, 000 cells, 500 genes for the linear process and 500 genes for the cyclic process. In the fourth example, we simulated 10, 500 cells, 400 genes for the cyclic process and 200 genes for each sub-lineage process. For all these samples, we adopted the following model parameters: *γ*_1_ = 25, *µ*_*d*_ = 0, *µ*_*α*_ = 0, *σ*_*d*_ = 0.25, *σ*_*α*_ = 0.25. We chose *T* = 10 in the fourth example while *T* = 15 in the other examples. We chose *γ*_2_ = 2 for simulating the cyclic process while *γ*_2_ = 8 for simulating the other processes. In simulation experiments, genes along a circular trajectory are ordered by their angular coordinates of the first two non-trivial Diffusion Map eigenvectors.

#### 8.2.2 Generalizing count model using negative binomial distribution to account for overdispersion

To investigate the impact of dispersion on the performance of GeneTrajectory, specifically in terms of gene ordering, we performed a negative binomial variant of our second and third simulation experiments in Figure 2. For each dataset, we simulated three distinct sparsity levels (5%, 10%, and 20%). For each sparsity level, we tested four different dispersion levels (parameterized by *θ*), each comprising 10 technical replicates. A lower *θ* value indicates higher dispersion. We evaluated the consistency between the inferred gene ordering and the ground truth by calculating their Spearman correlation (Supplementary Figure 1). It shows that GeneTrajectory exhibits remarkable stability across all sparsity and dispersion levels.

#### 8.2.3 Experimental details of mouse embryo skin sample preparation

##### Mice

K14Cre (Dassule et al., 2000) mice were bred to Wntless^*fl/fl*^ (Carpenter, 2010) mice. A random population of both male and female embryos were used for all experiments. All procedures involving animal subjects were performed under the approval of the Institutional Animal Care and Use Committee of the Yale School of Medicine.

##### EdU administration

To assess proliferation, EdU was administered to pregnant mice intraperitoneally (25 *µ*g/gm) and embryos were harvested after 1.5 hour.

##### In-situ hybridization

10% formalin-fixed paraffin embedded (FFPE) whole embryos were used for histological analysis. FFPE specimens were subsectioned at 10 µm thickness. The RNAscope Multiplex Fluorescent Detection Kit v2 (ACDBio, 323110) was used for single-molecule fluorescence in situ hybridization according to the manufacturer’s protocol. Briefly, subsections were deparaffinized, permeabilized with hydrogen peroxide followed by antigen retrieval and protease treatment before probe hybridization. After hybridization, amplification and probe detection was done using the Amp 1-3 reagents. Probe channels were targeted using the provided HRP-C1-3 reagents and TSA fluorophores: Cy3 (AkoyaBio, NEL744001KT), Cy5 (AkoyaBio, NEL745001KT), fluorescein (AkoyaBio, NEL741001KT). EdU staining was done using the Click-it EdU Imaging Kit Alexa 488 (Life technologies, c10338) according to manufacturer’s instructions. Nuclear counter-stain was done using Hoechst 33342 (Invitrogen, H3570) before mounting with SlowFade Mountant. RNA scope probes used (ACDBio): Mm-Lef1 (441861) and Mm-Sox2 (401041).

##### Microscopy

FISH paraffin-embedded images were acquired using the Leica TCS SP8 Gated STED 3X super-resolution confocal microscope with a 40x oil immersion (NA 1.3) objective lens, scanned at 5 *µ*m thickness, 1024×1024 pixel width, 400 Hz.

##### Single-cell dissociation

Embryonic dorsolateral/flank skin was micro-dissected from E14.5 littermate control and mutant embryos and dissociated into a single-cell suspension using 0.25% trypsin (Gibco, Life Technologies) for 20 minutes at 37*°C*. After genotyping, 2-3 embryos were pooled by condition. Single-cell suspensions were then stained with DAPI (Fisher Scientific, NBP2-31156) just prior to fluorescence-activated cell sorting.

##### Fluorescence-activated cell sorting

DAPI-excluded live skin cells were sorted on a BD FACS Aria II (Biosciences) sorter with a 100 *µ*m nozzle. Cells were sorted in bulk and submitted for 10X Genomics library preparation at 0.75-1.0×10^6^/mL concentration in 4% FCS/PBS solution.

##### H-score quantification

For quantification based on FISH, cells with 4-5 dots were considered positive (according to the RNAScope manufacturer’s instructions) and subsections from a total of n=4 different embryos were examined. To measure RNA expression levels, H-scores were calculated according to ACDBio manufacturer’s instructions: a cell with 0 dot is scored 0, 1-3 dots score 1, 4-9 dots score 2, 10-15 dots and/or less than 10% clustered dots score 3, and more than 15 dots and/or more than 10% clustered dots score 4; then the final H-score of a given cell type A is calculated by summing the (% cells scored B within all cells in A)*B for score B in 0-4.

##### Single-cell RNA sequencing and library preparation

Chromium Single cell 3’ GEM Library and Gel Bead Kit v3.1 (PN-1000121) was used according to the manufacturer’s instructions in the Chromium Single Cell 3’ Reagents Kits V3.1 User Guide. After cDNA libraries were created, they were subjected to Novaseq 6000 (Illumina) sequencing. For each scRNA-seq experiment, control and littermate mutant samples were prepared in parallel at the same time, pooled, and sequenced on the same lane.

#### 8.2.4 Analytical details of real-world examples

##### Human myeloid dataset analysis

Myeloid cells were extracted from a publicly available 10x scRNA-seq dataset (https://support.10xgenomics.com/single-cell-gene-expression/datasets/3.0.0/pbmc_10k_v3). QC was performed using the same workflow in (https://github.com/satijalab/Integration2019/blob/master/preprocessingscripts/pbmc_10k_v3.R). After standard normalization, highly-variable gene selection and scaling using the Seurat R package [56], we applied PCA and retained the top 30 principal components. Four sub-clusters of myeloid cells were identified based on Louvian clustering with a resolution of 0.3. Wilcoxon rank-sum test was employed to find cluster-specific gene markers for cell type annotation.

For gene trajectory inference, we first applied Diffusion Map on the cell PC embedding (using a local-adaptive kernel, each bandwidth is determined by the distance to its *k*-nearest neighbor, *k* = 10) to generate a spectral embedding of cells. We constructed a cell-cell *k*NN (*k* = 10) graph based on their coordinates of the top 5 non-trivial Diffusion Map eigenvectors. Among the top 2, 000 variable genes, genes expressed by 0.5% *−* 75% of cells were retained for pairwise gene-gene Wasserstein distance computation. The original cell graph was coarse-grained into a graph of size 1, 000. We then built a gene-gene graph where the affinity between genes is transformed from the Wasserstein distance using a Gaussian kernel (local-adaptive, *k* = 5). Diffusion Map was employed to visualize the embedding of gene graph. For trajectory identification, we used a series of time steps (11, 21, 8) to extract three gene trajectories. Gene ordering was done based on the algorithm described in 8.1.4.

##### Mouse embryo skin data analysis

We separated out dermal cell populations from the newly collected mouse embryo skin samples (8.2.3) (aligned to the mouse genome mm10 by CellRanger v6.1.2). Cells from the wildtype and the Wls mutant were pooled for analyses. After standard normalization, highly-variable gene selection and scaling using Seurat, we applied PCA and retained the top 30 principal components. Three dermal celltypes were stratified based on the expression of canonical dermal markers, including *Sox2, Dkk1*, and *Dkk2*. For gene trajectory inference, we first applied Diffusion Map on the cell PC embedding (using a local-adaptive kernel bandwidth, *k* = 10) to generate a spectral embedding of cells. We constructed a cell-cell *k*NN (*k* = 10) graph based on their coordinates of the top 10 non-trivial Diffusion Map eigenvectors. Among the top 2, 000 variable genes, genes expressed by 1% *−* 50% of cells were retained for pairwise gene-gene Wasserstein distance computation. The original cell graph was coarse-grained into a graph of size 1, 000. We then built a gene-gene graph where the affinity between genes is transformed from the Wasserstein distance using a Gaussian kernel (local-adaptive, *k* = 5). Diffusion Map was employed to visualize the embedding of gene graph. For trajectory identification, we used a series of time steps (9, 16, 5) to sequentially extract three gene trajectories. To compare the differences between the wiltype and the Wls mutant, we stratified Wnt-active UD cells into seven stages according to their expression profiles of the genes binned along the DC gene trajectory.

##### Cell-cycle (CC) gene trajectory validation

We extracted the Cyclebase [57] gene list from the Supplementary Table 5 in [58], in which genes are categorized into groups of G1/S, S, G2, G2/M, and M phase markers. We also incorporated histone genes into the S phase gene list as they are upregulated during the S phase for the active synthesis of histone proteins [59]. We plotted the distribution of genes from different phases along the gene trajectory associated with the CC process in the dermal example (Extended Data Fig. 3b). We observed that genes corresponding to the G1/S phase were located around the start of the gene trajectory, followed by a group of genes highly expressed during the S phase. G2M-related genes were located along the second half of the gene trajectory. Specifically, G2 genes appeared in the middle of the trajectory, followed by a group of genes regulating the switch from G2 to M. Genes associated with the M phase were found around the end of the trajectory. This indicates that GeneTrajectory can effectively capture gene dynamics associated with different phases of the cell cycle.

##### Different visualizations of gene embedding

Gene embedding visualization is agnostic to gene-gene distance computation and trajectory identification. Different ways of gene embedding visualization for the two real-world examples included in the manuscript are shown and compared in Supplementary Figure 2. We would advise users to apply diffusion-based visualization techniques, e.g., diffusion map or PHATE [60], to display the trajectories, as they were designed to capture and reveal the connectivity of graphs.

##### Assessing the stability of capturing gene processes in the dermal example

After identifying three prominent gene trajectories by running GeneTrajectory on the original cell graph (with the maximum of iteration = 5000 when calculating gene-gene distances), we constructed a new cell graph using only the genes extracted from each gene trajectory. We then reran the gene trajectory inference on each new cell graph for 1) all the genes, and 2) the same set of genes that were used to construct the new cell graph (Supplementary Figure 3). We found that the ordering of the genes used to define the new cell graph stayed in a high degree of consistency with their original ordering inferred by our method (when we constructed the cell graph using all genes). This consistency highlights the stability of GeneTrajectory in inferring gene dynamics underlying each process, unaffected by the presence of coexisting gene programs and biological effects.

Meanwhile, we observed potential caveats of iteratively running GeneTrajectory on the cell graphs constructed using the genes along a previously identified gene trajectory. This is because, in each iteration, the cell graph is only determined by the subset of genes corresponding to a specific process. There is no theoretical guarantee that the cell graph still encodes the geometric information necessary for identifying a gene trajectory associated with a different process. In other words, the new cell graph may distort the cell geometry for the other processes.

#### 8.2.5 Comparing Wasserstein metric to other canonical metrics for learning gene geometry

We conducted an extensive benchmark on using different metrics (including the Earth-Mover distance, Euclidean distance, Pearson correlation distance, Spearman correlation distance, Cosine similarity, total variation distance (equivalent to L1 distance or Manhattan distance in its discrete form), Jensen-Shannon distance, and Hellinger distance) to learn gene geometry in simulation datasets (Supplementary Figure 4). Datasets for evaluation were generated based on simulations (corresponding to the second and third simulation examples in Fig. 2). Specifically, we simulated datasets with three different sequencing depths (i.e., the percentage of non-zero entries in the gene-by-cell count matrix = 2.5%, 5%, 10%), each having 10 replicates. To evaluate the performance, we calculated the Spearman correlation between each inferred gene ordering and the ground truth. The Wasserstein distance recovers gene ordering more accurately and robustly than other metrics.

#### 8.2.6 Comparing GeneTrajectory with cell trajectory methods in terms of gene ordering inference

We performed a benchmark to compare GeneTrajectory with five representative cell trajectory inference methods, Monocle2 [16], Monocle3 [10], Slingshot [9], PAGA [11], and CellRank [15]. We assessed their performances on two types of datasets:

- simulation datasets (corresponding to third simulation example in Fig. 2) with varying sparsity levels of the count matrix (i.e., the percentage of non-zero entries in the gene-by-cell count matrix = 2.5%, 5%, 10%, 20%) and different numbers of cells (500, 1, 000, 2, 500).
- the real-world dermal dataset depicted in Fig. 4-5 with or without cell-cycle-effects regression.

For these cell trajectory inference approaches, after cell pseudotime inference, we leveraged generalized additive models (GAM) using the mgcv [61] (Mixed GAM Computation Vehicle with Automatic Smoothness Estimation) R package to smooth the gene expression along the cell pseudotime, followed by ordering the genes based on their peak locations. For the simulation datasets, we calculated the Spearman correlation between the true gene order and the inferred gene order by each method. For the dermal dataset, the assessment is done by examining the ordering of experimentally-verified markers during dermal condensate (DC) differentiation.

In summary, Monocle2 and Monocle3 require the specification of a starting (root) cell state to generate cell pseudoorder. In simulation experiments, we chose the cell with the ground truth pseudotime *t* = 0 as the starting cell state. In the dermal dataset, we first looked at the diffusion map embedding of cells to define the tip cell (expressing Sox2) as the terminal cell state of DC differentiation process. We then chose the upper dermal cell that has the largest distance (in the transcriptome space) to the terminal cell state as the starting cell state. PAGA and SlingShot require specification of a starting cell cluster to create cell pseudotime. Based on the same strategy as described above, we chose the cluster containing the starting cell state as the staring cell cluster. The core steps in each analysis workflow for cell trajectory inference methods are summarized below:

- SlingShot: We used the getLineages function to construct the minimum spannning tree(s) on cell clusters. We then fitted principal curves using the getCurves function, which served as the basis for cell pseudotime construction.
- PAGA: Cells were reclustered using the leiden method implemented in the Scanpy toolkit. We constructed the PAGA graph of these cell clusters, and inferred progression of cells through geodesic distance along the graph using scanpy.tl.dpt.
- Monocle2: We employed the built-in DDRTree method for cell dimension reduction. We used the orderCells function to generate the cell ordering while the root state was defined by the starting cell state as noted above.
- Monocle3: Cells were partitioned using the built-in Louvain method. We learned the principal graph across all partitions and then ordered the cells using the order cells function.
- CellRank: For the simulation experiments, since we don’t have the information about the spliced and unspliced read counts, we used CellRank’s CytoTRACEKernel to infer the transition dynamics and cell pseudotime. For the dermal example, we applied CellRank based on RNA velocity inference. Specifically, the spliced/unspliced counts were quantified by the velocyto toolkit. We used scVelo’s dynamical model [62] to infer RNA velocities. CellRank was then applied to infer the initial states & terminal states of transition and construct cell lineages. We selected the cell lineage that terminates its transition at the DC cell population, and fitted GAM models (built in CellRank) to order the genes along the cell pseudotime of the selected lineage.

#### 8.2.7 Hyperparameter selection guidelines and robustness evaluation

We would advise users to choose and determine the parameters according to the following standards:

- If users choose to use diffusion maps (or PCA) to generate a cell embedding. The number of eigenvectors (or principal components) for cell graph dimensionality reduction can be ascertained by examining the eigenvalues in descending order to identify an eigen-gap or the point where the spectrum starts to flatten out.
- K in cell kNN-graph construction is a user-defined hyperparameter.The chosen value for K should ensure the cell graph is fully connected.
- The number of gene programs is determined by the number of branches (gene trajectories) identifiable from the gene graph. This determination is made interactively during the process of branch identification. Specifically, when a new branch is being extracted, we exclude the genes that have already been assigned to existing branches. Subsequently, we identify one of the remaining genes that is most distant from the origin of diffusion embedding as the tip of the next branch. If the remaining genes visually form an indistinct cloud that does not exhibit a trajectory structure, we cease the process of branch identification.
- The time step *t* for random walks in each iteration of branch identification is interactively determined by inspecting the gene embedding. Specifically, when *t* increases, a greater number of genes are incorporated as the members of the branch to be extracted. The optimal *t* for extracting each branch should yield the longest trajectory without incorporating the genes in the indistinct cloud.
- The number of gene bins for visualization is determined by the resolution users wish to inspect for shifting patterns in gene distributions over cell embedding. An ideal number would be between 5-10. The choice of bin number does not affect gene trajectory inference.

We conducted an extensive evaluation to assess the robustness of GeneTrajectory with varying combinations of parameters. These parameters included *K* for constructing cell-cell kNN graphs, *Ndim* for dimensionality reduction, and *K*_*a*_ for determining local adaptive kernel bandwidths in diffusion map construction. To assess GeneTrajectory’s performance on simulated datasets, we computed the Spearman correlation between the inferred gene ordering and the ground truth ordering. For the real-world examples, we performed a cross-validation by examining the Spearman correlation between all pairs of inferred gene orderings. The results of this evaluation are depicted in Supplementary Figures 5-7. Specifically, these experiments include:

- We simulated bifurcation datasets and cylindrical datasets (corresponding to the second and third simulation examples in Fig. 2) with varying sparisity levels (i.e., the percentage of non-zero entries in the gene-by-cell count matrix = 2.5%, 5%, 10%, 20%, each has 10 replicates; each replicate includes 1, 000 cells). We tested GeneTrajectory using a combination of *K* = 5, 10, 15, 20, 25 and *Ndim* = 5, 10, 15, 20, 25. The evaluation outputs are shown in Supplementary Figure 5.
- Using the same simulation datasets above, we tested GeneTrajectory using *K*_*a*_ = 5, 10, 15, 20, 25 for constructing the diffusion embedding of cells. The evaluation outputs are shown in Supplementary Figure 6.
- In two real-world examples included in this manuscript, we tested GeneTrajectory on cell graphs constructed using different numbers of eigenvectors (*Ndim* = 5, 10, 15, 20, 25). The evaluation outputs are shown in Supplementary Figure 7.

## 9 Data Availability

The human peripheral blood mononuclear cell (PBMC) scRNA-seq dataset is available at https://support.10xgenomics.com/single-cell-gene-expression/datasets/3.0.0/pbmc_10k_v3. The mouse embryonic skin dataset generated and analyzed in this study is available from the Gene Expression Omnibus (GEO) with the accession number GSE255534. The processed Seurat data objects for these two datasets are available at Figshare (dx.doi.org/10.6084/m9.figshare.25243225). The Cyclebase gene list was extracted from the Supplementary Table 5 in [58].

## 10 Code Availability

The R package of GeneTrajectory and code used for data analysis will be available on GitHub.

---

**Extended Data Figure 1.**
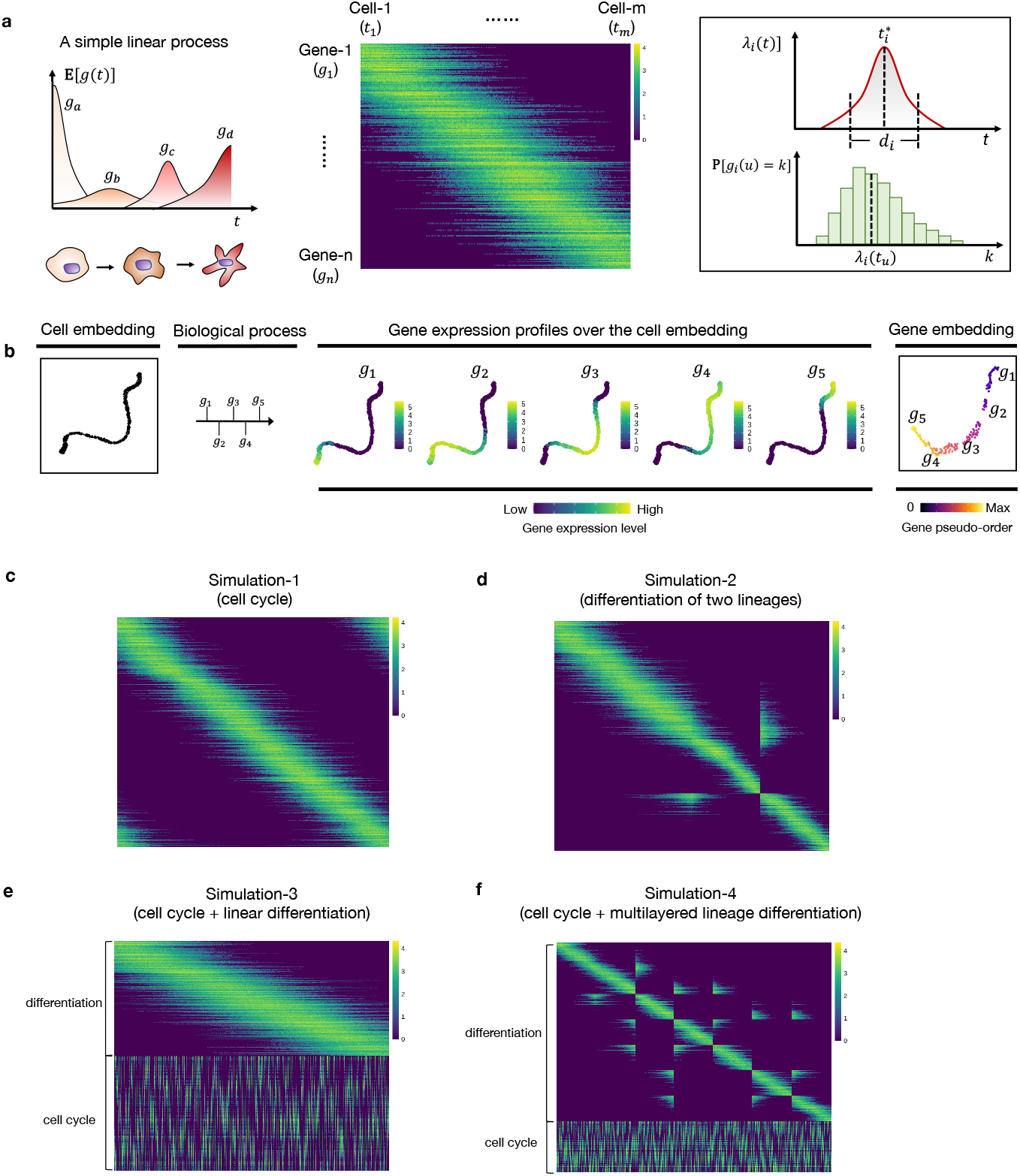

**Extended Data Figure 2.**
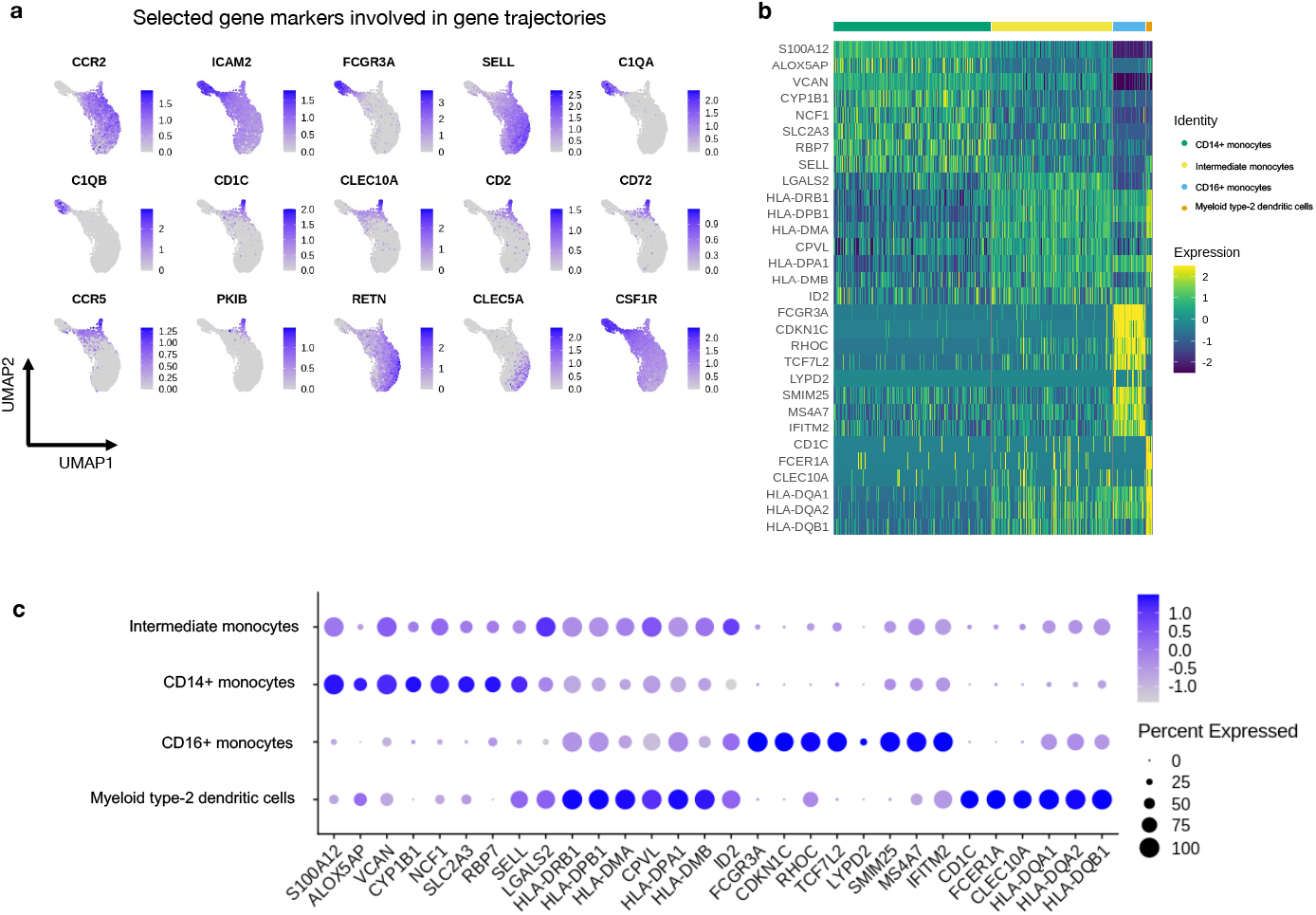

**Extended Data Figure 3.**
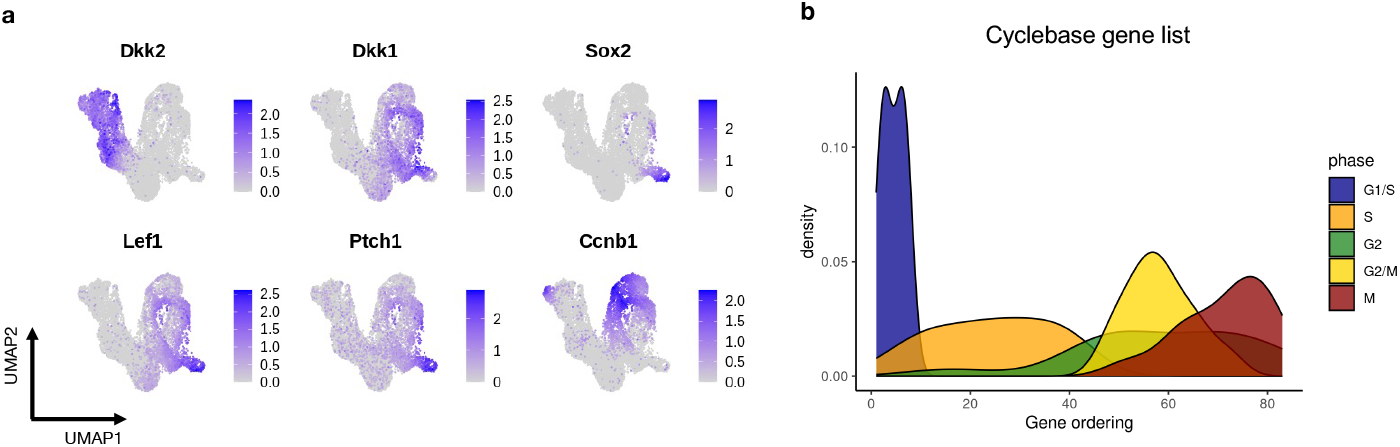

**Extended Data Figure 4.**
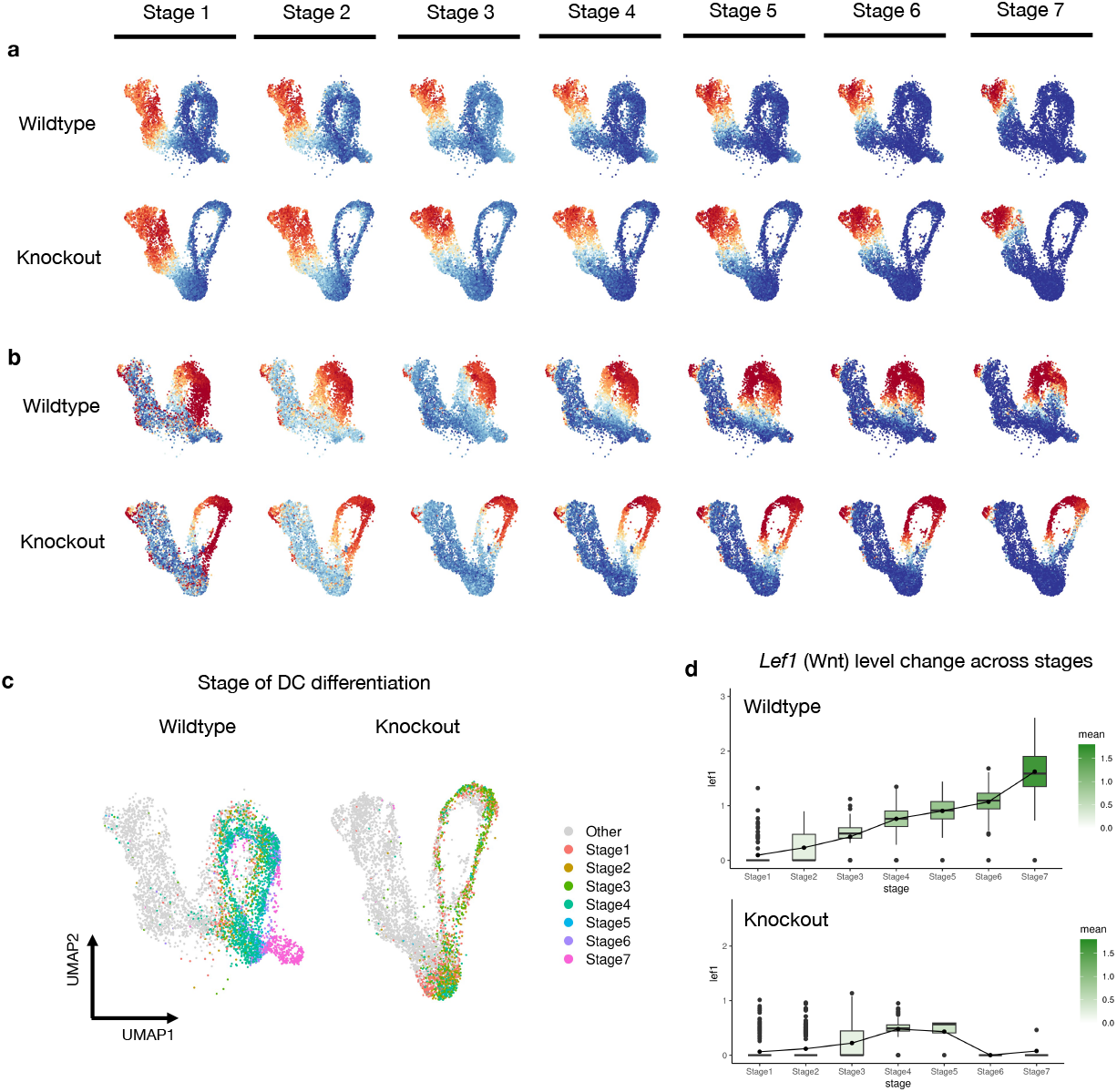

**Extended Data Figure 5.**
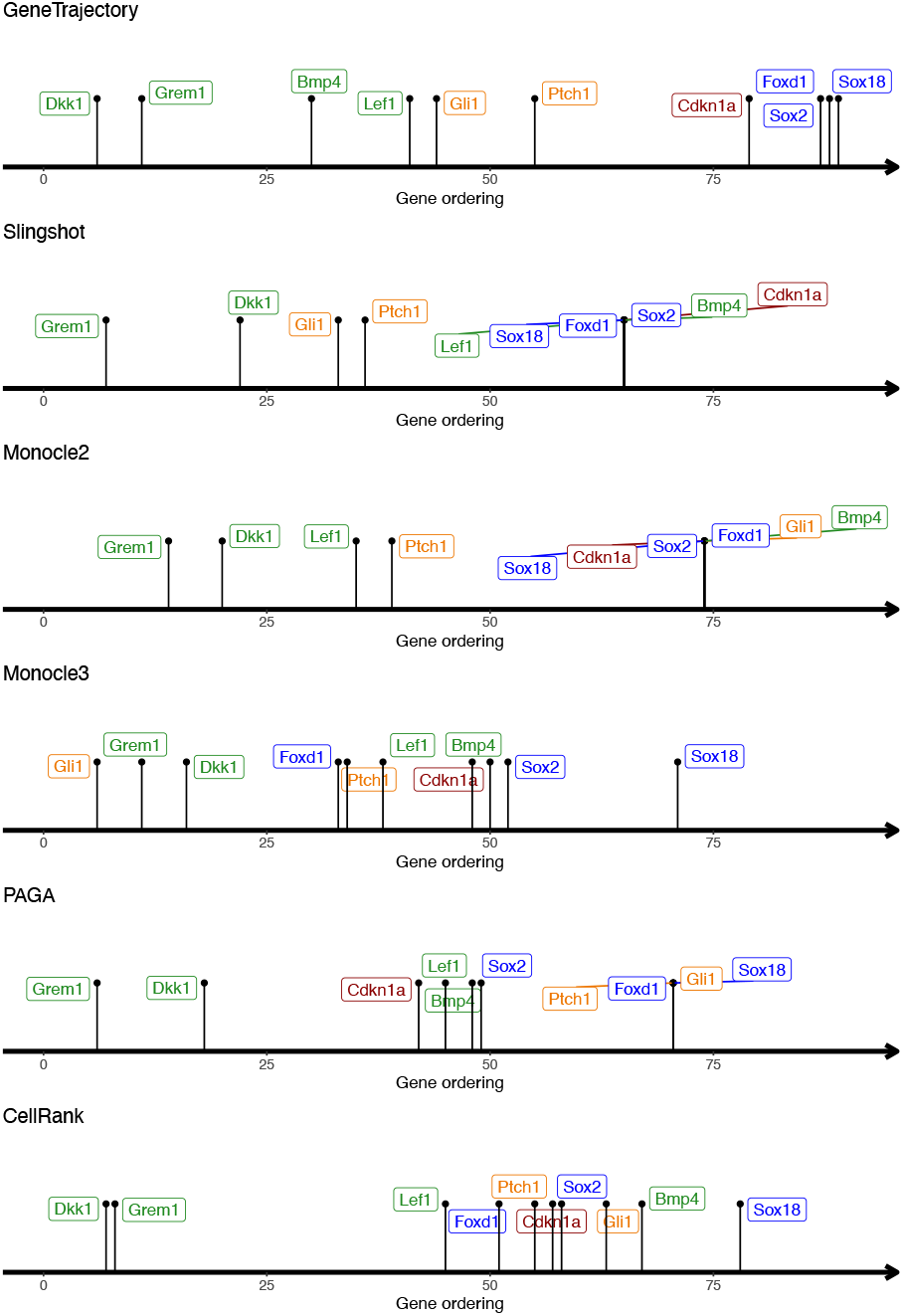

## Notes

### Summary of Updates

The manuscript and supplementary materials have been revised.

