## Supplementary Figures for "Gene Trajectory Inference for Single-cell Data by Optimal Transport Metrics"

Supplementary Figure 1

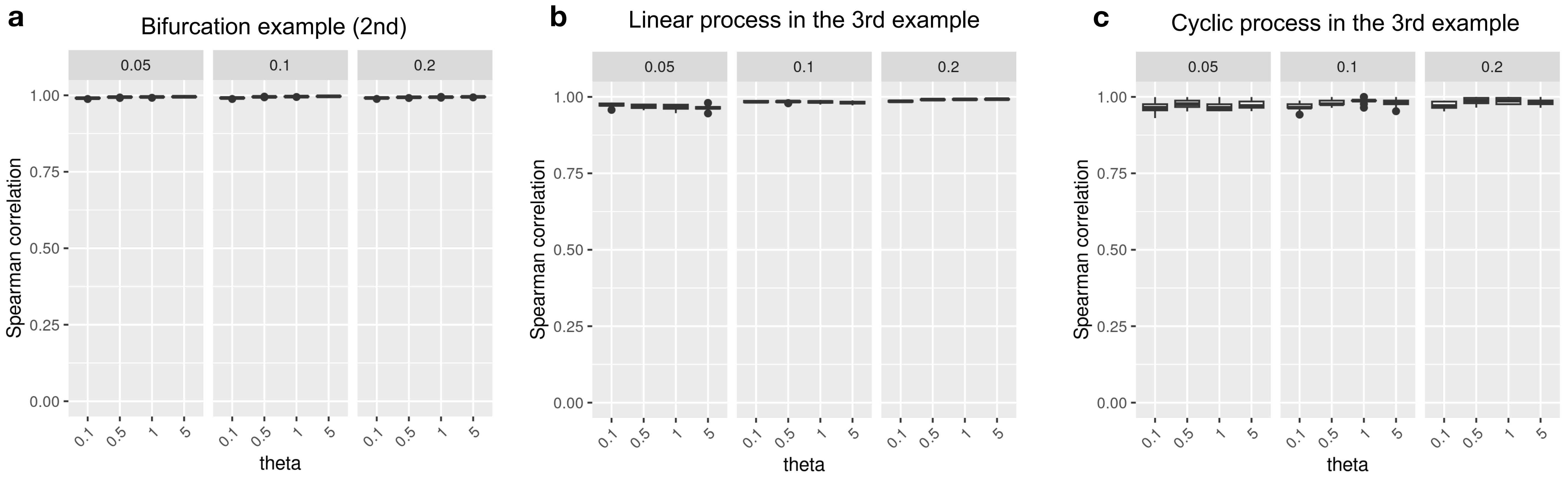


**Supplementary Figure 1 | GeneTrajectory ordering is stable with respect to variable dispersion in negative binomial simulations.**

**a,** the bifurcation process from the second simulation example of Fig. 2;

**b,** the linear process from the third simulation example of Fig. 2;

**c,** the cyclic process from the third simulation example of Fig. 2.

Each panel corresponds to a different sparsity level of the count matrix, as indicated in the header. Within each panel, each column represents a specific dispersion level of the count matrix (parameterized by theta), each comprising 10 technical replicates. The Y-axis denotes the Spearman correlation between the inferred gene ordering and the ground truth.

Box plots: The box represents the interquartile range (IQR), with the line inside the box indicating the median. Whiskers extend to a maximum of 1.5 × IQR beyond the box, with outliers represented as individual points.

### Supplementary Figure 2

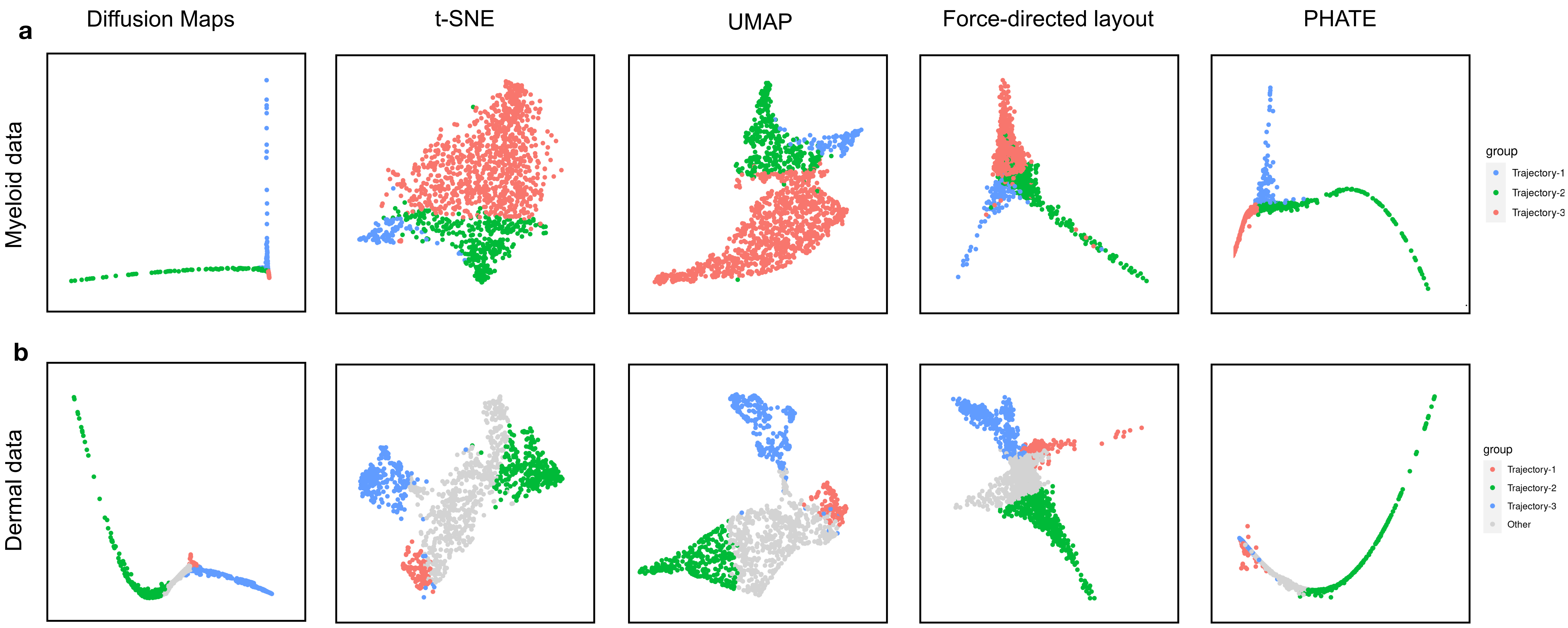


**Supplementary Figure 2 | Different visualizations of gene geometry**

Different visualizations (including Diffusion Maps, t-SNE, UMAP, force-directed layout, and PHATE) showing the gene embeddings of (**a**) the myeloid example and (**b**) the dermal example.

Supplementary Figure 3

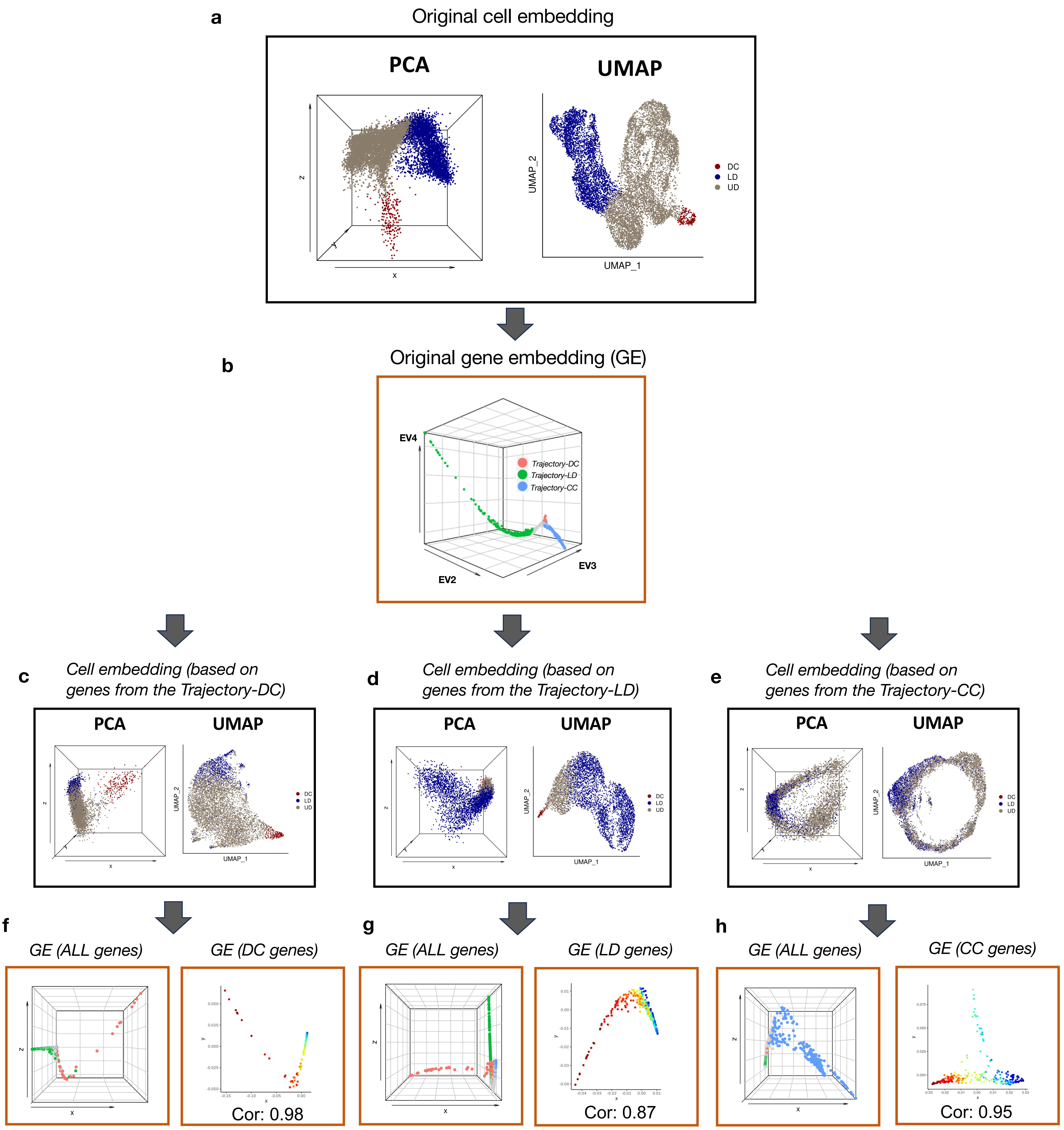


**Supplementary Figure 3 | Evaluating the stability of GeneTrajectory in inferring gene ordering for each process**

**a**, PCA and UMAP cell embeddings of the dermal data (corresponding to Fig. 4-5).

**b**, DM embedding of genes (using the leading three non-trivial EVs) showing the gene trajectories.

**c-e**, PCA and UMAP cell embeddings constructed by only using genes along the DC, LD, and CC gene trajectories.

**f-h**, DM (using the leading three non-trivial EVs) showing the gene embedding (GE) of 1) all genes (color-coded by the initially identified gene trajectories), and 2) the same set of genes that are used to construct each new cell graph (colored based on their initially inferred orderings).

Supplementary Figure 4

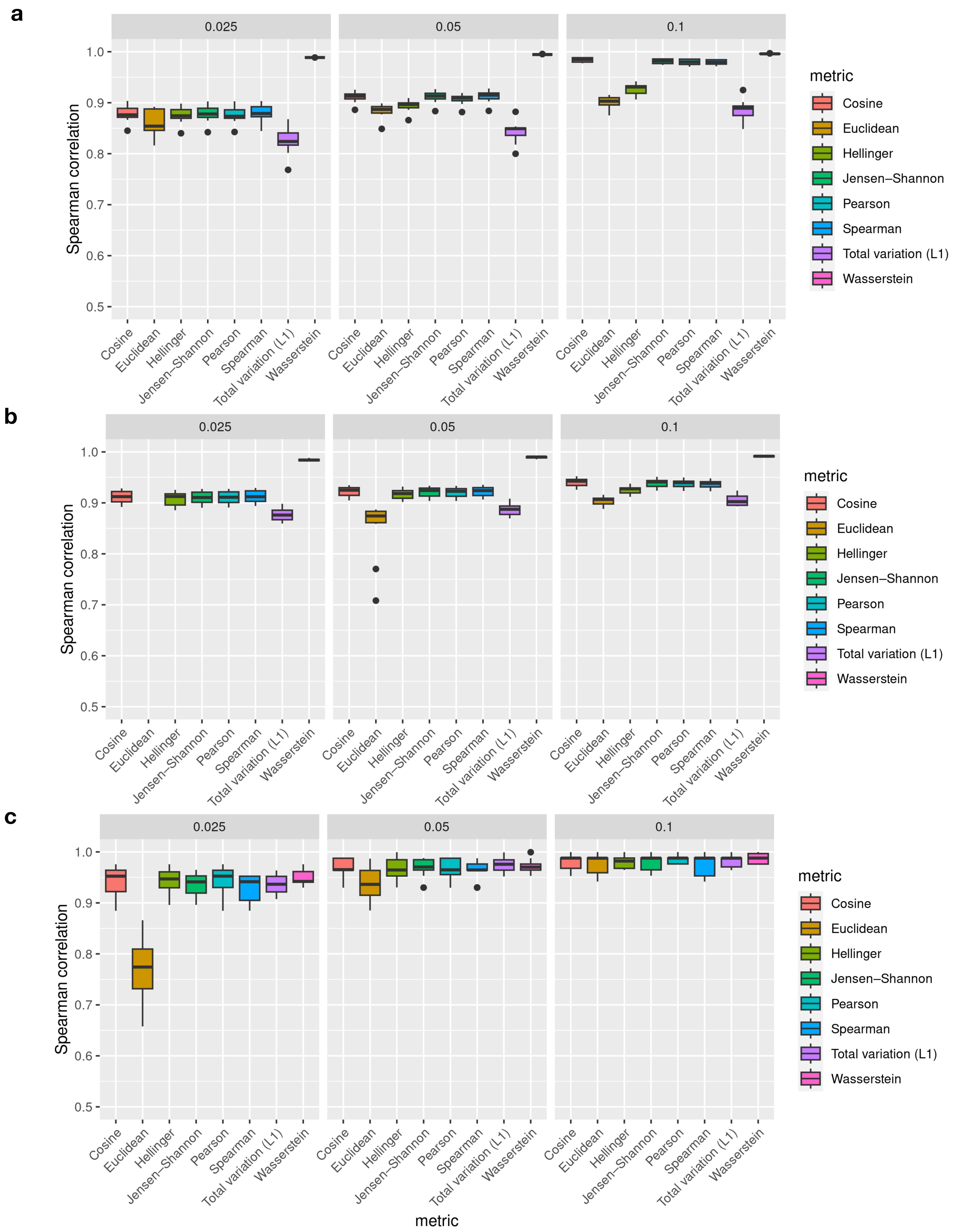


**Supplementary Figure 4 | Wasserstein distances outperform many alternative distance metrics for gene-gene embedding.**

Benchmark experiments on using different gene-gene distance metrics for gene ordering inference along

**a**, the bifurcation process from the second simulation example of Fig. 2;

**b**, the linear process from the third simulation example of Fig. 2;

**c**, the cyclic process from the third simulation example of Fig. 2.

Each panel corresponds to a different sparsity level of the count matrix, as indicated in the header.

Box plots: The box represents the interquartile range (IQR), with the line inside the box indicating the median. Whiskers extend to a maximum of 1.5 × IQR beyond the box, with outliers represented as individual points.

### Supplementary Figure 5

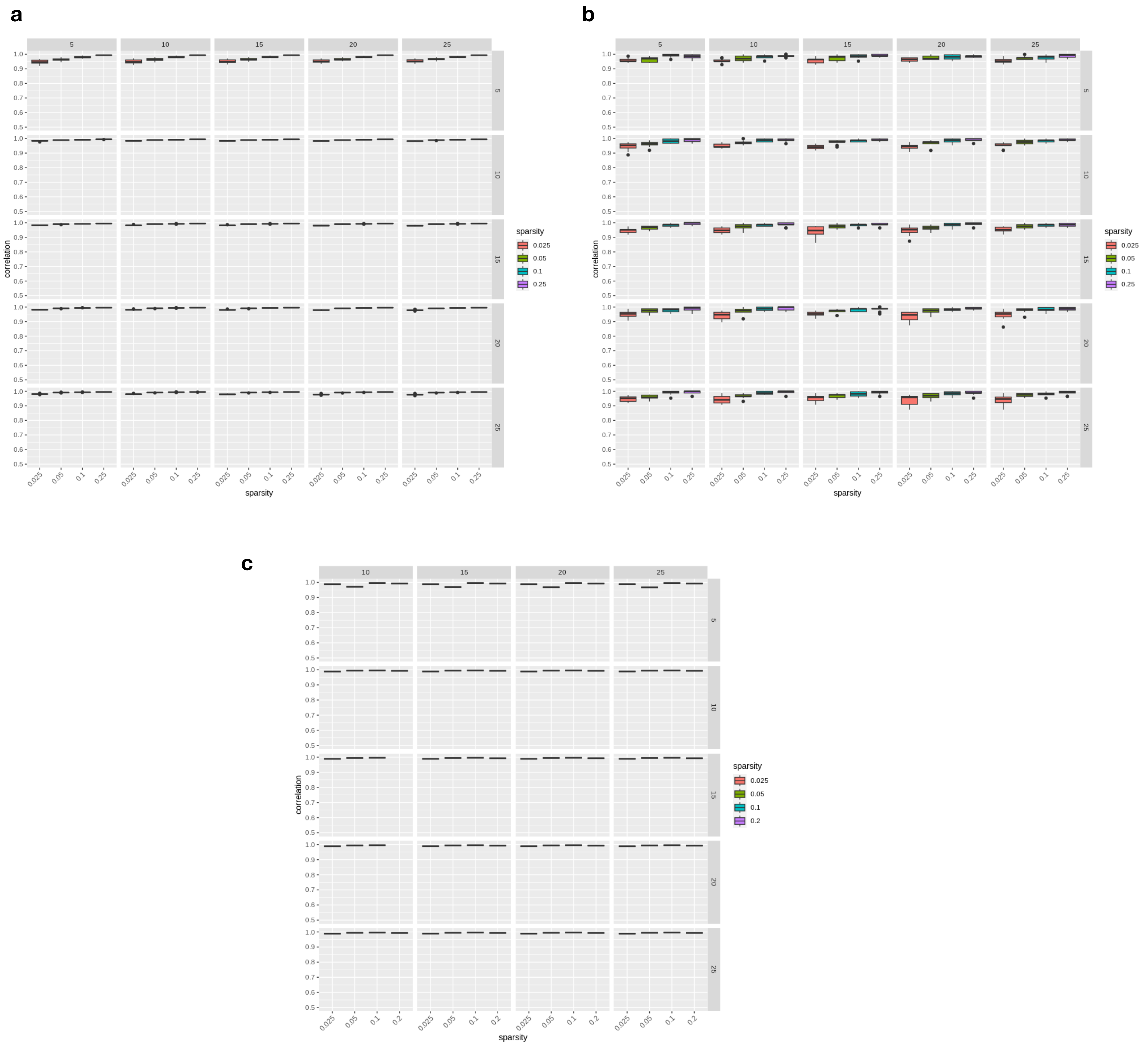


**Supplementary Figure 5 | Hyperparameter robustness evaluation of gene ordering inference**

**a**, Evaluating the gene ordering along the linear process in the third simulation example of Fig. 2;

**b,** Evaluating the gene ordering along the cyclic process in the third simulation example of Fig. 2;

**c**, Evaluating the gene ordering along the bifurcation process (corresponding to the second simulation example in Fig. 2).

Each row in the table corresponds to a specific value of Ndim (the number of leading non-trivial eigenvectors used to build the cell graph), while each column corresponds to a specific value of K (for defining the cell kNN graph). For every unique combination of Ndim and K, the performance is evaluated on datasets with four distinct sequencing depths, each having 10 replicates. The Missing values in section **c** are due to the disconnectedness of the cell graph when using the corresponding combination of hyperparameters.

Box plots: The box represents the interquartile range (IQR), with the line inside the box indicating the median. Whiskers extend to a maximum of 1.5 × IQR beyond the box, with outliers represented as individual points.

Supplementary Figure 6

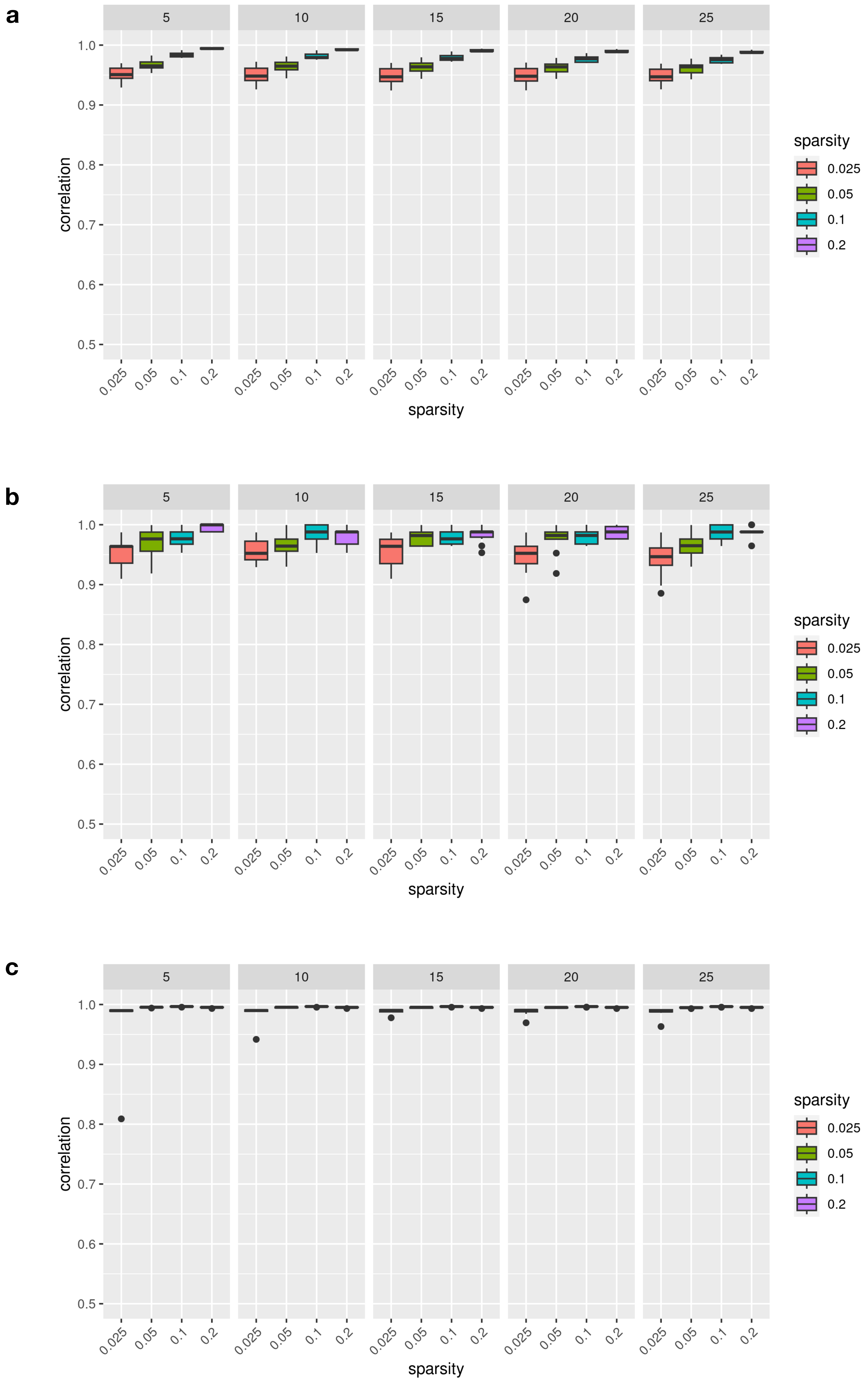


**Supplementary Figure 6 | Robustness evaluation of gene ordering inference on the kernel bandwidth $K_a$ for constructing the diffusion embedding of cells**

**a**, Evaluating the gene ordering along the linear process in the third simulation example of Fig. 2;

**b,** Evaluating the gene ordering along the cyclic process in the third simulation example of Fig. 2;

**c**, Evaluating the gene ordering along the bifurcation process (corresponding to the second simulation example in Fig. 2).

Each panel corresponds to a specific $K_a$ as labeled in the header. For each $K_a$, performance on datasets with four different sequencing depths (each having 10 replicates) is assessed.

Box plots: The box represents the interquartile range (IQR), with the line inside the box indicating the median. Whiskers extend to a maximum of 1.5 × IQR beyond the box, with outliers represented as individual points.

Supplementary Figure 7

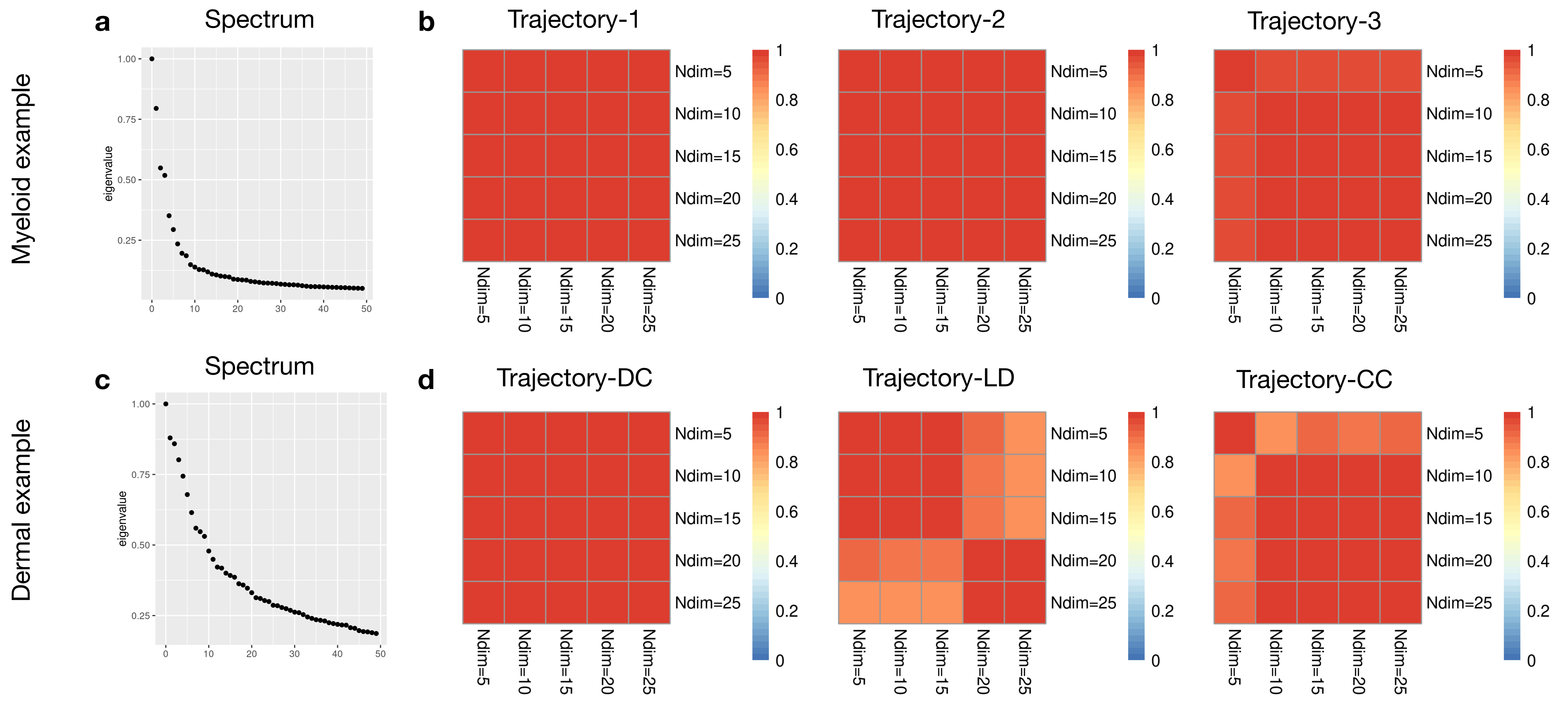


**Supplementary Figure 7 | Robustness evaluation of gene ordering inference based on Ndim (the number of eigenvectors for constructing the cell embedding) in real-world biological datasets**

**a-b**, Evaluation on the myeloid example (Fig. 3);

**c-d**, Evaluation on the dermal example (Fig. 4-5).

**a, c,** Spectrums of the random walk affinity matrices, showing the eigenvalues in a descending order to identify an eigen-gap or the point where the spectrum starts to flatten out.

**b, d,** Heatmaps showing the Spearman correlations between the gene orderings inferred by using different numbers of eigenvectors to construct the cell graph.
